# Spatial and temporal profiling of the complement system uncovered novel functions of the alternative complement pathway in brain development

**DOI:** 10.1101/2023.11.22.568325

**Authors:** Yingying Zhang, Brianna Watson, Ajitanuj Rattan, Tyrone Lee, Smriti Chawla, Ludwig Geistlinger, Yilin Guan, Minghe Ma, Barbara J. Caldarone, Wenchao Song, Jeffrey R. Moffitt, Michael C. Carroll

## Abstract

Mounting evidence implicated the classical complement pathway (CP) in normal brain development, and the pathogenesis of neuropsychiatric and neurodegenerative diseases. However the source and regulation of complement in the brain remain unclear. Using MERFISH, a spatial transcriptomic method with single-cell resolution, we established a developmental brain atlas of the complement system. We showed that the brain synthesizes essential building blocks of the complement system locally with remarkable cellular and spatial heterogeneity. We provided transcriptional evidence supporting the presence of the alternative pathway (AP), but lack of lectin pathway (LP) activity in postnatal brain development. Cell type, temporal and spatial expression patterns of genes involved indicate non-redundant functions of the CP and AP. In addition, deficiency in Masp3-driven AP resulted in developmental and cognitive defects, indicating essential functions of the AP, an observation that highlights the necessity to disentangle differential involvement of the three complement activation pathways in development and diseases.

## Introduction

The complement system was first identified over a century ago and is a well characterized key regulator of both peripheral innate and adaptive immunity (Carroll, 2004; Ricklin et al., 2010). There are over 50 known complement related genes in rodents, consisting of core complement components, receptors and regulators. Complement cascade can be activated through three distinct pathways: the classical pathway (CP), the lectin pathway (LP) and the alternative pathway (AP). There have been numerous studies showing the classical complement pathway has been co-opted in the nervous system to mediate developmental synapse pruning. During this process, C1q, the first member to activate the classical complement pathway, tags excessive synapses to be eliminated, and recruits and activates C4 and C3 at the synaptic terminal. Such decorated synapses can then be recognized by the complement receptor Cr3 expressed on the microglia surface and subsequently engulfed by microglia (Presumey et al., 2017; Schafer et al., 2012; Stephan et al., 2012; Stevens et al., 2007). Recent genetic and animal model studies have shown that over activation of the complement cascade can cause Schizophrenia, possibly through excessive synaptic pruning (Druart et al., 2021; Schizophrenia Working Group of the Psychiatric Genomics, 2014; Sekar et al., 2016; Yilmaz et al., 2021). Moreover, complement activation has also been frequently implicated in the progression of neurodegenerative diseases (Dalakas et al., 2020), including Alzherimer’s disease (Hong et al., 2016; Krance et al., 2021; Shi et al., 2017), Huntington’s disease (Wilton et al., 2023), amyotrophic lateral sclerosis (ALS) (Kjaeldgaard et al., 2018), and multiple sclerosis (Watkins et al., 2016; Werneburg et al., 2020). However, unlike in the periphery, our understanding of the complement system in the brain remains incomplete. For example, the expression and regulation of the complement cascade in different brain regions, cell types, and across development have not been comprehensively determined. And, while the role of the CP in neuronal development is clear, it remains unknown if the LP and AP are involved in brain development, and if so, whether their functions are redundant with the CP.

Systemic profiling of the complement cascade gene expression and regulation in the central nervous system has significant challenges. First, it is a complex system that encompasses over 50 genes. Second, many complement genes are lowly expressed or are confined to specific cell populations in the brain. These properties make the complement system difficult to be characterized by traditional single-cell sequencing methods due to constraints in sequencing depth, cell capture efficiency, and relatively low throughput in the number of cells being profiled. In addition, the intricate heterogeneity of the brain complicates matters, as spatial variability in complement gene expression is not captured by conventional sequencing methods based on dissociated cells, yet such spatial variability could have important functional implications. MERFISH (Multiplexed Error-Robust Fluorescent *in situ* Hybridization), a spatial transcriptomic method with single-cell resolution, has emerged as an effective solution (Chen et al., 2015; Moffitt et al., 2018; Moffitt et al., 2016) as it has the demonstrated ability to profile hundreds to thousands of genes, including lowly expressed genes, and from these measurements define, discover, and map cell types in different regions of the mouse and human brain (Allen et al., 2023; Fang et al., 2022; Zhang et al., 2021). Recently, MERFISH and related methods have been extended to define and map cell populations across the entire adult mouse brain (Shi et al., 2023; Yao et al., 2023). By then integrating these measurements with companion single-cell RNA sequencing datasets, the transcriptome-wide gene expression and spatial organization of thousands of neuronal populations has been described. However, only very limited number of genes in the complement pathway were included in the set of genes targeted in these studies. Given the limited sensitivity of single-cell RNA sequencing, inferred complement gene expression through data integration remains low in resolution and accuracy. Moreover, these studies have been limited to adult mice, leaving the developmental regulation, a critical aspect of complement function in the brain, unexplored.

We report here a comprehensive spatial atlas of the complement system in mouse brains over postnatal development. We found that 47 out of 51 complement genes examined are locally expressed in postnatal mouse brains. Expression of many complement genes are developmentally regulated, and exhibited remarkable heterogeneity in cell type and spatial expression patterns. These measurements shed light on the role of the AP in brain development. Specifically, the serine proteases essential to activate the LP were not expressed anywhere in the brain, suggesting the absence of the LP during normal postnatal brain development. By contrast, our data unveiled a distinctive, developmentally and spatially regulated expression pattern in Masp3, a serine protease key to the AP activation (Gullipalli et al., 2023; Hayashi et al., 2019), suggesting an unappreciated role for the AP in neuronal development. Masp3 deficiency caused reduced birth rate, lower body weight and cognitive deficits, supporting essential developmental functions of Masp3-driven AP. In addition, cell type, spatial and temporal expression patterns of AP components are inversely correlated with those for CP, indicating divergent functions of the two complement activation pathways in brain development.

Together our study provided a high resolution spatial and developmental overview of the complement system expression and regulation in the brain. Furthermore, our data unveiled an unexpected role of the AP in the brain, and therefore emphasize the necessity to distinguish respective roles of the three complement activation pathways in development and disease pathogenesis.

## Results

### Complement components are locally produced in the brain with cell-type and brain-region heterogeneity

To construct a brain atlas for the complement system, we designed a 249-gene MERFISH probe library comprising genes in the following groups: 1) canonical markers for major brain cell types; 2) cortical layer and brain region markers; 3) neurotransmitter synthesis, transport and receptors; 4) known genes involved in neural development and homeostasis; and 5) 51 complement genes, including complement components, receptors and regulators (Table S1). Given the tremendous heterogeneity of cell types in the brain, and the lack of consensus of terminal neuronal and glial cell types, our goal with this library was to determine complement expression in major categories of cells in the brain parenchyma.

To maximize the capture of spatial heterogeneity, we performed MERFISH on six whole sagittal sections taken from two distinct locations of 60-day-old (p60) adult mouse brains: medial sections (1000±100 μm from the midline, n=3 mice) and lateral sections (2450±100 μm from the midline, n=3 mice). These two locations were chosen to cover various thalamic nuclei and to capture brain region-to-region variabilities. Using an established MERFISH data processing pipeline (Moffitt et al., 2018), we decoded over 250 million RNA molecules that were assigned to a total of 513,520 segmented cells. Overall, the measured gene expression was highly reproducible between replicates (Figure S1A) and well correlated with published abundance values from brain RNA-sequencing (Figure S1B). In addition, the spatial distribution of RNAs included to mark different brain regions detected by MERFISH matched those from the Allen Brain Atlas (ABA) *in-situ* hybridization database (Figure S1C).

Using this library, we identified 16 major brain cell types based on established cell type markers (Figures 1A-1B, S1D), including two types of excitatory neurons (vGlut1^+^ and vGlut2^+^ respectively), inhibitory neurons, dopaminergic neurons in the substantia nigra, and cholinergic neurons in the medulla. For non-neuronal cells, we identified astrocytes, microglia, oligodendrocytes, oligodendrocyte progenitor cells, pericytes, endothelial cells, ependymal cells and cells of choroid plexus. While our goal was not to define all neuronal types, we, nonetheless, performed secondary clustering and identified refined cell clusters (Figures 1D-1K, S2), which resulted in 17 subtypes of vGlut1^+^ neurons, 12 subtypes of vGlut2^+^ neurons, and 12 subtypes of inhibitory neurons. When projected to MERFISH coordinates, vGlut1^+^ neurons mainly occupied the cortex and cerebellum, and its subclusters formed distinct cortical layers and sub-structures of the hippocampus that are characteristic of the anatomical organization of the cerebral cortex (Figures 1D, S2A-S2B). vGlut2^+^ neurons mainly located in the brain stem, and segregated into various nuclei in the thalamus and mid-/hind- brain (Figures 1E, S2C-S2D). Inhibitory neurons formed transcriptionally distinct subclusters in the cortex, striatum, mid-/hind-brain and the cerebellum. In the thalamus however, inhibitory neurons were densely located the reticular nucleus (IN-RT), but sparsely distributed in other thalamic nuclei (Figures 1F, S2E-S2F). Among the non-neuronal cell types, astrocytes (Figures 1G, S2G-S2H) and oligodendrocytes (Figures 1H, S2I-S2J) showed spatial heterogeneity. Cortical (PP2) and brainstem (PP1) protoplasmic astrocytes were transcriptionally distinct from each other (Figure 1G), with brainstem astrocytes marked by expression of GABA transporters Slc6a1/GAT1 and Slc6a11/GAT3, and cortical astrocytes marked by Mfge8 and Mertk (Figure S2H). Gfap positive fibrous astrocytes and mature oligodendrocytes (Figures 1H, S2I-S2J) are concentrated along the fiber tracts, notably the corpus callosum. On the other hand, microglia (Figures 1I, S2K-S2L), oligodendrocyte progenitor cells (Figures 1J, S2M-S2N), and endothelial cells (Figures 1K, S2O-S2P) didn’t display brain-region specific sub-populations using the genes included in our library. Leveraging distinct spatial distribution patterns of neuronal subtypes, we assigned 23 distinct brain regions to each slice (Figure 1C).

**Figure 1:**
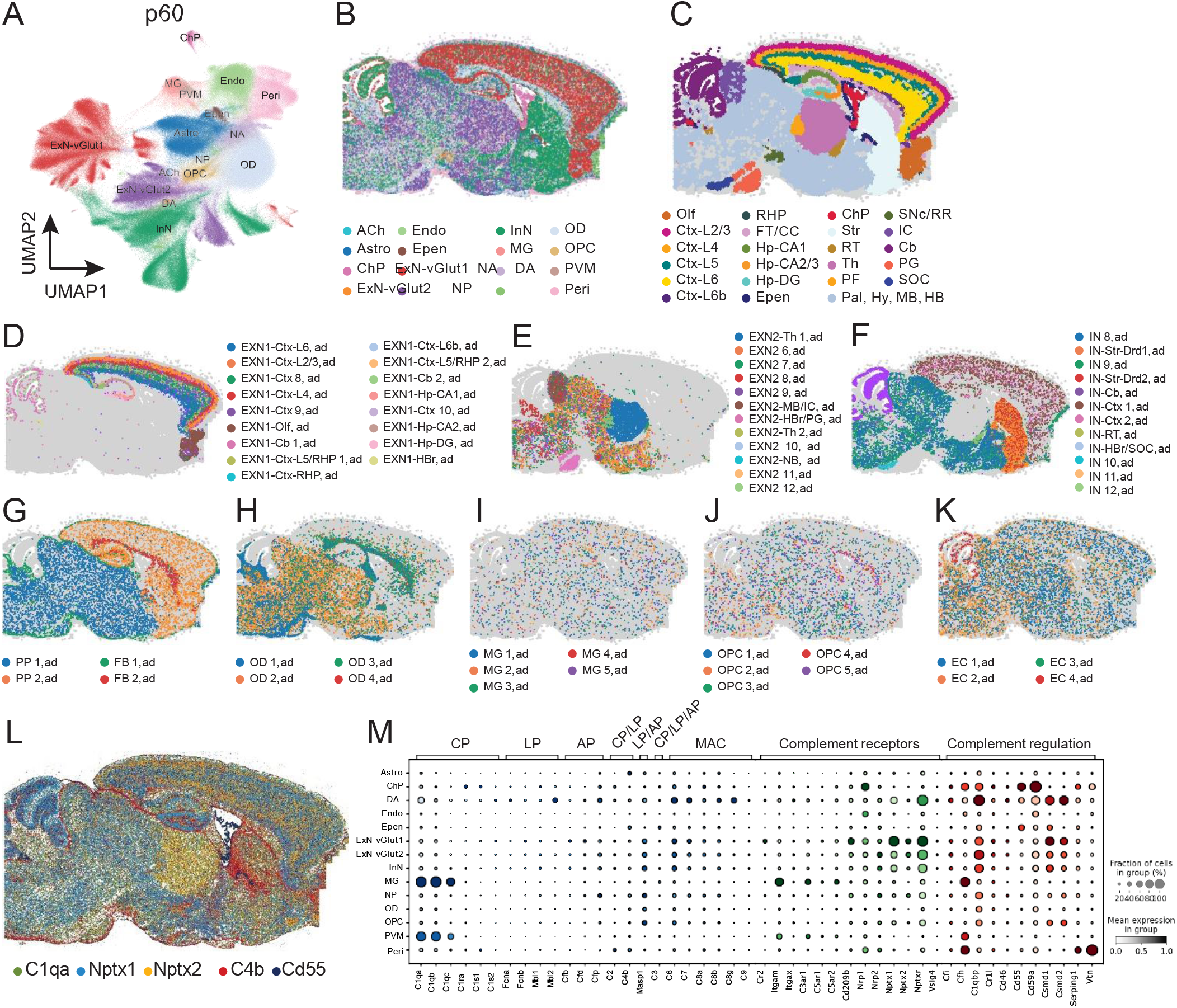
Complement expression in adult mouse brain. **(A)** UMAP visualization of primary cell clusters identified by MERFISH in p60 mouse brain sagittal sections. **(B)** Representative plot of spatial localization of each primary cell type. Cells are colored by cell type as in (A). **(C)** Representative plot of brain regions identified by MERFISH. Cells are colored by brain region. **(D-K)** Spatial localization of secondary cell clusters of vGlut1^+^ (ExN-vGlut1, D) and vGlut2^+^ (ExN-vGlut2, E) excitatory neurons, inhibitory neurons (InN, F), astrocytes (Astro, G), oligodendrocytes (OD, H), microglia (MG, I), oligodendrocyte progenitor cells (OPC, J) and endothelial cells (Endo, K). Lists of cell type and brain region abbreviations are provided in supplementary tables 7 and 8 respectively. **(L)** Representative plot of spatial distribution of transcripts of C1qa (green), Nptx1 (light blue), Nptx2 (orange), C4b (red) and Cd55 (dark blue) detected in a p60 mouse sagittal brain slice MERFISH. **(M)** Dot plot of complement expression by cell type. Color intensity corresponds to average gene expression, dot size corresponds to the percentage of cells expressing each gene.

To evaluate overall expression level and spatial heterogeneity of complement genes, we first examined RNA spatial distribution of each individual complement gene and noticed vast heterogeneity in overall abundance and spatial distribution among them. C1qa, a well-studied pattern recognition molecule activating the CP, is widely expressed and evenly distributed throughout the brain. In contrast, expression of C4b that functions downstream of C1qa, is concentrated at the brain border and surrounding the lateral ventricle. Neuronal pentraxins Nptx1 and Nptx2 are newly identified brain specific complement C1q receptors (Kovacs et al., 2020; Zhou et al., 2023). Nptx1 is highly expressed throughout the brain except in the thalamus and striatum, whereas Nptx2 is mainly expressed in the cortex, thalamus and striatum. Expression of a subset of complement genes are more spatially restricted. For example, Cd55, or decay accelerating factor (DAF) that inactivates C3 and C5 convertase (Lublin and Atkinson, 1989; Medof et al., 1984), is exclusively expressed at the choroid plexus (Figure 1L). The heterogeneity in both spatial distribution and relative abundance of complement genes met our expectation based on published single cell RNA sequencing studies, and further emphasize the necessity of using a highly sensitive spatial method covering the entire mouse brain to profile the complement pathway in the brain.

We next evaluated the cell type expression pattern of the complement genes. As false positive signals are present in image-based approaches to single-cell transcriptomics, such as MERFISH, we have included 8 barcodes that were not assigned to any target RNA in the library (termed blanks) to measure this false positive rate. We then used the per-cluster expression of these blank barcodes to determine genes with expression values greater than these blanks (two standard deviations above the average of blanks). 44 out of 51 (86.3%) complement genes examined were expressed above this threshold in at least one cluster. C1rb, Hc (or C5), C9, C4bp, Fcna, Masp2 and Vsig4 did not pass this threshold in adult samples (Table S2).

Among the genes involved in CP, the C1q pattern recognition complex (C1qa, C1qb, C1qc) was highly and exclusively expressed by microglia, consistent with findings in published single cell RNA sequencing studies in both human and mouse brains (Fonseca et al., 2017; Herring et al., 2022; Zeisel et al., 2018). Although the C1q complex has been frequently studied in the context of complement activation in the brain, C1r and C1s, serine proteases essential to drive CP activation, are commonly overlooked. Our data showed that in general, CP serine proteases, C1ra, C1s1, C1s2, were lowly and sparsely expressed, indicating restricted baseline CP activity in the brain. C1ra and C1s1, were mainly expressed by the choroid plexus and pericytes (Figure 1M), implying that CP serine proteases could be made available through circulating CSF produced by the choroid plexus and through paracrine secretion by pericytes that are found in the base membrane of capillaries. As for LP, its pattern recognition molecules (Fcna, Fcnb, Mbl1, and Mbl2) were much less abundant than their counterparts in the CP, and the critical serine protease of LP, Masp2, was below detection limit. Although there have been reports indicating the LP activity in prenatal neuronal migration (Gorelik et al., 2017a) and neurodegenerative conditions (Kjaeldgaard et al., 2018; Mercurio et al., 2020), it is likely inactive in adult brain at steady state. AP is initiated by low-grade C3 hydrolysis. Factor D (Cfd) facilitates proteolytic activation of factor B (Cfb). Activated factor B drives the formation C3 convertase C3bBb and the amplification of the pathway. Properdin (Cfp) is considered a positive regulator of the AP by stabilizing C3bBb (Hourcade, 2006). All components for AP activation can be detected. Neurons are top cell types that express AP components Cfd, Cfb and Cfp, and notably neuronal progenitors expressed the highest level of Cfp. In contrast to the CP, cellular expression pattern of AP genes indicates a neuronal cell autonomous regulation of the AP activation, and a possible role in neuronal progenitor cells.

Complement receptors are also integral components of the complement cascade. Microglia functions as brain tissue resident macrophages, and similar to their peripheral counterparts, they express canonical complement receptor Itgam (or Cd11b/Cr3), anaphylatoxin receptors C3ar1, C5ar1 and C5ar2, which mediate microglia activation and phagocytic functions. Conversely, neurons, notably vGlut1^+^ excitatory neurons, synthesize neuronal pentraxins Nptx1, Nptx2 and Nptxr that function as C1q receptors and serve as docking site for the CP activation (Kovacs et al., 2020; Zhou et al., 2023).

All three complement activation pathways converge at the proteolytic activation step that converts C3 to C3b, which in the periphery, further triggers the formation of membrane attack complex (MAC), known as the terminal step of the complement cascade. MAC formation process starts with the conversion of C5 (or Hc) to generate C5b, which in turn recruits C6, C7 and C8 consecutively. Eventually multiple units of C9 are recruited and form a pore on the target cell plasma membrane. Formation of MAC can be catastrophic and may lead to lytic cell death, and therefore is tightly regulated. In the periphery, although C5-C9 are constitutively expressed primarily in the liver and secreted to the circulation, multiple regulatory proteins prohibit the formation of MAC, including soluble complement regulators vitronectin (or complement S-protein) and clusterin (Sheehan et al., 1995; Tschopp et al., 1993), and membrane bound regulators Cd55/DAF (Lublin and Atkinson, 1989; Medof et al., 1984) and Cd59 (known as the MAC inhibitory protein) (Rollins and Sims, 1990). The choroid plexus, a blood-cerebral-spinal fluid (CSF) barrier tissue that is deeply imbedded in the brain ventricles, is highly vascularized with fenestrated capillaries (Dani et al., 2021; Solar et al., 2020), and is in contact with abundant source of complement components found in the blood. We detected remarkably high levels of membrane bound MAC regulatory protein Cd55 and Cd59a expression in the choroid plexus (Figure 1M), necessary to protect it from complement attack. Unlike peripheral tissues, at steady state, an intact blood-brain barrier protects the brain parenchyma from exposure to complement components in the blood. In our data, local expression of C5 and C9 were below detection limit (Table S2), indicating transcription-level suppression of MAC formation in the brain parenchyma. To provide further protection against MAC, neurons expressed membrane bound CP inhibitor Csmd1 (Escudero-Esparza et al., 2013; Kraus et al., 2006) (Figure 1M). In addition, pericytes and microglia expressed additional soluble complement regulatory proteins including vitronectin, C1 inhibitor Serping1 and AP inhibitor complement factor H (Cfh) (Ferreira et al., 2010; Gorelik et al., 2017b; Sheehan et al., 1995). These additional protections are of importance especially under pathological conditions involving blood-brain barrier damage.

Collectively, we showed that the brain has the structural basis for complement activation with aspects that are distinct from the periphery. First, transcriptional data support the presence of CP and AP, but the lack of LP in healthy brain, whereas in the periphery, LP provides antibody independent first line defense against pathogenic microorganism infection (Degn et al., 2011). In addition, although C1q complex is highly expressed by microglia, expression of C1s, C1r, serine proteases essential for the CP activation, are very limited in the brain, suggesting minimal basal level CP activation. And finally, MAC formation is likely suppressed under steady state due to lack of local production of C5 and C9, preventing the brain cells from lytic cell death. These brain-specific aspects provide transcriptional regulation of complement activation. Further complement regulation is assured upon expression of soluble complement regulatory proteins by microglia, pericytes and the choroid plexus, and neuronal expression of membrane bound complement regulatory proteins.

### Developmentally regulated complement expression revealed a novel role of Masp3 in early postnatal cortical development

Complement activation has been shown to regulate postnatal synaptic pruning (Schafer et al., 2012; Stephan et al., 2012; Stevens et al., 2007), a highly dynamic process that happens at different ages depending on the brain region. It is conceivable that complement expression patterns not only vary spatially, but also change over development. Thus, a comparison of the temporal expression of complement cascade would be informative for identifying critical periods of different complement components in brain development, and distinguishing developmental versus homeostatic function of the complement system. We therefore performed MERFISH using the same target gene library on six additional sagittal sections (3 medial sections 960±80µm from the midline, and 3 lateral sections 2040±80µm from the midline, n=3 mice) from three 5-day-old (p5) newborn mouse brains. p5 and p60 samples were integrated by normalizing average total RNA counts per cell. Due to remarkable differential expression related to development, we were concerned that integrating datasets from the two age groups would mask biological variability and skew cell type identification. Thus, we clustered the p5 and p60 samples separately. We identified major cell clusters in p5 samples similar to those in adult samples, except for mature oligodendrocytes and cholinergic neurons (Figures 2A-2B, S3A). Sub-clustering of major cell types further identified 14 subtypes of vGlut1^+^ excitatory neurons that segregated by cortical layers and hippocampal regions (Figures S3B, S4A-S4B), 15 subtypes of vGlut2^+^ excitatory neurons that grouped into distinct nuclei and regions in the brain stem (Figures S3C, S4C-S4D), and 10 subtypes of inhibitory neurons that also showed distinct brain region specificity (Figures S3D, S4E-S4F). Leveraging the spatial distribution patterns of neuronal subclusters, 22 distinct brain regions were assigned to p5 samples (Figure 2C). Comparing overall transcript abundance between the p5 and p60 samples, neurogenesis and proliferation markers such as Mki67, Dcx, Sox11, and Cd24a were significantly enriched in p5 samples, as expected. In addition, a subset of neurotransmission genes were also among the top differentially expressed genes between the two ages (Figure 2D).

**Figure 2:**
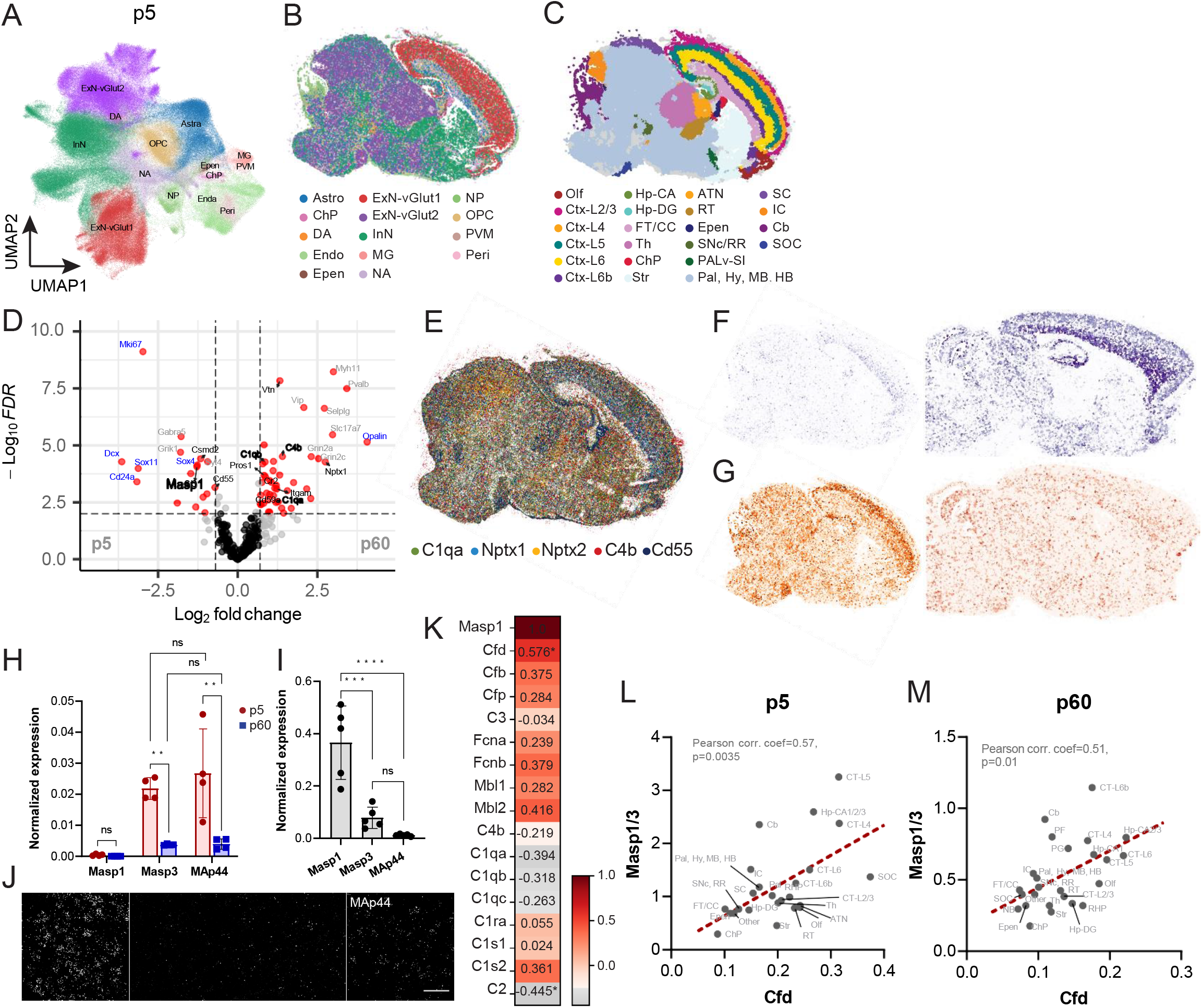
Masp3 is a developmentally regulated serine protease of the alternative complement pathway in the brain. **(A)** UMAP visualization of primary cell clusters identified by MERFISH in p5 mouse brain sagittal sections. **(B)** Representative plot of spatial localization of each primary cell type. Cells are colored by cell type as in (A). **(C)** Representative plot of brain regions identified by MERFISH. Cells are colored by brain region. Lists of cell type and brain region abbreviations are provided in supplementary tables 9 and 8 respectively. **(D)** Volcano plot showing differentially expressed genes in p5 vs. p60 mouse brains detected by MERFISH. All slices and all cell types are included in the analysis. Genes with over 1.6-fold difference between p5 and p60, and with false discovery rate (FDR, Calculated by Benjamini-Hochberg test for multiple comparisons) less than 0.01 are highlighted in red. **(E)** Representative plot of spatial distribution of transcripts of C1qa (green), Nptx1 (light blue), Nptx2 (orange), C4b (red) and Cd55 (dark blue) detected in a p5 mouse sagittal brain slice by MERFISH. **(F, G)** Representative plot of transcripts of Nptx1 (F) and Masp1/3 (G) detected by MERFISH in p5 (left panel) and p60 (right panel). **(H)** Expression of Masp1, Masp3 and MAp44 in p5 and p60 brains by ddPCR. n=4 mice per age group; Two-way ANOVA with Sidak’s multiple comparisons test; error bars represent standard deviation; ns: not significant; **: p<0.01. **(I)** Masp1, Masp3 and MAp44 expression in adult mouse liver by ddPCR. n=4 mice. One-way ANOVA with Tukey’s multiple comparisons test; error bars represent standard deviation; ns: not significant; ns: not significant; ***: p<0.001; ****: p<0.0001. **(J)** Representative image of RNAscope with isoform non-specific probe for Masp1/3 (pan-Masp1/3, i), isoform specific probe for Masp1 (ii), Masp3 (iii), and MAp44 (iv) in adult mouse parafascicular region. **(K)** Heatmap of Pearson correlation coefficient of Masp1/3 expression by brain region with other complement components as detected by MERFISH in p5 mouse samples. Pearson correlation coefficient ρ is labeled on the heatmap. Complement components below detection threshold in both age groups were not included in the analysis. *: p<0.05. **(L, M)** Scatter plot of average normalized counts per cell of Cfd (X-axis) and Masp1/3 (Y-axis) by brain region detected by MERFISH in p5 (L) and p60 (M) brains, showing significant positive correlation between the spatial expression distribution of the two genes.

Overall, complement expression in early postnatal mouse brains exhibited a comparable cell type distribution as seen at p60 (Figure S2M). In p5 mouse brains, seven complement genes, C1ra, C1rb, C1s1, Hc, C4bp, Cr2 and Masp2 were below detection limit (Table S3), four of which, C1rb, Hc, C4bp and Masp2, were also below threshold in adult mouse brains. Therefore, 47 out 51 (92.2%) complement genes examined in MERFISH were expressed in the brain in at least one age group. However, there were noticeable differences in the spatial distribution of at least a subset of complement genes between the two age groups (Figure 2E, spatial distribution of C1qa, Nptx1, Nptx2, C4b and Cd55 RNA were plotted). In addition, differential gene expression analyses showed that throughout the brain, the expression of 15 complement genes were developmentally regulated (Table S4, FDR<0.05). Specifically, genes in the CP, C1qa, C1qb, C1ra, C4b are among the top enriched genes in adult mouse brains (Figure 2D). Consistently, complement receptor Itgam/Cd11b was 2.3 times more enriched (Table S4) and neuronal C1q receptor Nptx1 was 6.7 times more enriched in p60 compared to p5 brains (Figures 2D-2F, Table S4), indicating a general induction of the CP over development.

On the contrary, Masp1/3 (MBL associated serine protease 1/3) was the most downregulated complement gene in adult mouse brains (reduced by 2.4 folds, Figures 2D-2G, Table S4). Neuronal cell-type specific (Figures S5A-S5C), and brain region specific (Figures S5D-S5G) analyses showed heterogeneous developmental regulation of Masp1/3. The most prominent down-regulation of Masp1/3 expression was detected in vGlut1^+^ excitatory neurons (reduced by over 4 folds in p60 compared to p5 vGlut1^+^ neurons, Figure S5A), followed by a 2.2-fold reduction in p60 vGlut2^+^ excitatory neurons (Figure S5B). Masp1/3 level in inhibitory neurons was not developmentally regulated (Figure S5C), mainly because of relatively low level of expression in p5 inhibitory neurons (Figure S3I). Brain region specific analyses showed significant reduction of Masp1/3 in the cortex, thalamus and the cerebellum (Figures S5D, S5E, S5G respectively), but no significant difference in the striatum, a region that is enriched for inhibitory neurons (Figure S5F).

Masp1/3 is a complex gene. It encodes three functionally distinct splice variants: serine protease Masp1 that activates the LP; MAp44 that acts as a competitive lectin pathway inhibitor; and serine protease Masp3 that activates the AP in Cfd dependent manner (Degn et al., 2010; Hayashi et al., 2019). MERFISH probes for Masp1/3 were not designed to distinguish isoforms due to high sequence overlap. Thus, to compare the relative abundance of each Masp1/3 isoforms, we designed isoform specific primers and performed digital droplet PCR (ddPCR) on mouse brain and liver samples. Masp1 was not detected in the brain at p5 or p60. On the contrary, Masp3 and MAp44 can both be detected and expressed at a significantly higher level in p5 than in p60 samples (Figure 2H). The brain Masp1/3 isoform abundance was in clear contrast to that in the liver, where Masp1 was the most abundant isoform, followed by Masp3 and then MAp44 (Figure 2I). The relative abundance of Masp1/3 splice variants in the brain were also confirmed by RNAscope using custom designed isoform specific probes (Figure 2J, i: panMasp1/3, isoform non-specific; ii: Masp1; iii: Masp3; iv: MAp44). The contrasting relative isoform abundance of the Masp1/3 gene indicates unique complement regulation in the brain compared to the periphery. These expression patterns further argue for the lack of LP activity in postnatal mouse brains, which is supported by the observation that both Masp1 and Masp2, serine proteases indispensable for the LP activation, were below detection limit, and abundant level of the LP inhibitor MAp44. Conversely, the presence of Masp3 suggests possible AP activity in the brain, especially in developing brain and neural progenitor cells.

In addition to Masp3, expression of complement factors Cfd and Cfb is necessary for the AP activation. As they are secreted factors, we reasoned if the AP is active in the brain, we should observe spatial co-expression of these factors. We further hypothesized that the spatial co-expression analysis could be used to infer if AP and CP share redundant functions in the brain. Specifically, an inverse spatial correlation pattern, or no association, would indicate non redundant functions. To test this idea, we calculated the average expression per brain region, and average expression per cell type for complement genes and computed pairwise Pearson correlation. As our data suggest lack of LP activity, we focused on components of the CP and AP that were detected above threshold. In p5 mice, we observed positive correlation of spatial distribution among key CP components (C1qa, C1qb, C1qc, C2, C4b), and AP (Masp1/3, Cfd, Cfb, Cfp) components (Figure S6A). In addition, spatial distribution of members of the CP (C1qa, C1qb, C1qc, C2, C4b) appeared to be inversely correlated with those of the AP (Masp1/3, Cfb, Cfd). A similar observation was made in p60 samples as well (Figure S6C). However, when analyzing co-expression pattern by cell type, components in different complement activation pathways did not segregate, except for the C1q complex that is composed of C1qa, C1qb and C1qc (Figure S6B for p5 samples, and Figure S6D for p60 samples). These observations support the idea that spatial close proximity of complement component expression correlates with their biological function. Furthermore, an inverse correlation between CP and AP components implied divergent functions of the two complement activation pathways in the brain.

To address whether Masp3 expression in the brain indicates AP activity, we focused on the spatial co-expression analysis of Masp1/3 in particular with other complement components using MERFISH data. Among all the complement components detected above threshold, complement factor D (or Cfd), the direct downstream target of Masp3 to drive AP activation (Gullipalli et al., 2023; Hayashi et al., 2019), was the only complement component that showed significant spatial correlation with Masp1/3 at both ages (Pearson correlation coefficient ρ=0.57, p=0.0035 in p5, Figure 2K, 2L; ρ=0.51, p=0.01 in p60, Figure 2M). This observation supports the idea that expression of Masp1/3 detected by MERFISH is likely an indication of Masp3-dependent AP activation in the brain, rather than the LP or other complement-independent functions of Masp1/3.

### Masp3 expression in adult mice is concentrated in neurons in the parafascicular and pontine gray nuclei

As Masp1/3 emerged as an intriguing complement gene in the brain, we more closely examined its cellular and spatial expression patterns. Among the inhibitory neurons, Masp1/3 was most highly expressed by cortical somatostatin positive interneurons (IN-CT 2) and cerebellar inhibitory neurons (IN-Cb), but expressed at a much lower level in striatal neurons (Figures 3A-3B). Among the vGlut1^+^ excitatory neurons in the cortex, Masp1/3 expression exhibited cortical layer heterogeneity, and was most enriched in cortical layer 6b excitatory neurons, followed by cortical layer 6 and 4 (EXN1-CT-L6b, -L6, -L4 respectively). In comparison, its expression was much lower in cortical layers 2/3 and 5 (EXN1-CT-L2/3, -L5) (Figures 3C-3D). vGlut2^+^ excitatory neurons can be broadly divided into those found in the thalamus, and those found in the mid/hind brain regions. The thalamic excitatory neurons can be further divided into the parafascicular region (EXN2-Th2) and the rest of the thalamus (EXN2-Th1) (Figure 1E). In the thalamus, Masp1/3 expression segregated with the vGlut2^+^ excitatory neuron subclusters. It is only highly expressed in the parafascicular region (PF) but much less so in the rest of the thalamus (Figures 3E-3F). Among the mid/hind brain region excitatory neurons, the pontine gray nucleus (PG) stood out as another distinct brain region highly enriched for Masp1/3 expression (Figures 3E-3F). We validate the MERFISH observation of Masp1/3 spatial distribution in the thalamus by RNAscope using Lypd6b as a PF region marker (Figure 3G i). Not only did we confirm a clear enrichment of Masp1/3 in the PF region compared to the rest of the thalamus, it showed significant heterogeneity within the PF region, with the dorsal lateral side of the PF region expressed the highest levels of Masp1/3 (Figure 3G ii), and the ventral side with much diminished expression (Figure 3G iii), that is comparable to the neighboring region in the thalamus (Figure 3G iv, quantified in Figure 3H). Similar observation was also seen using an RNAscope probe specific to the Masp3 isoform (Figure 3I). The PF region can be further divided into molecularly distinct sub regions, each characterized by unique neuronal projection patterns and regulating specific aspects of motor behavior (Mandelbaum et al., 2019; Zhang et al., 2022). Therefore, spatial expression patterns of Masp3 suggest it may regulate motor activity and, function that has been associated with the lateral side of the PF region (Zhang et al., 2022) and the PG nuclei (Kratochwil et al., 2017).

**Figure 3:**
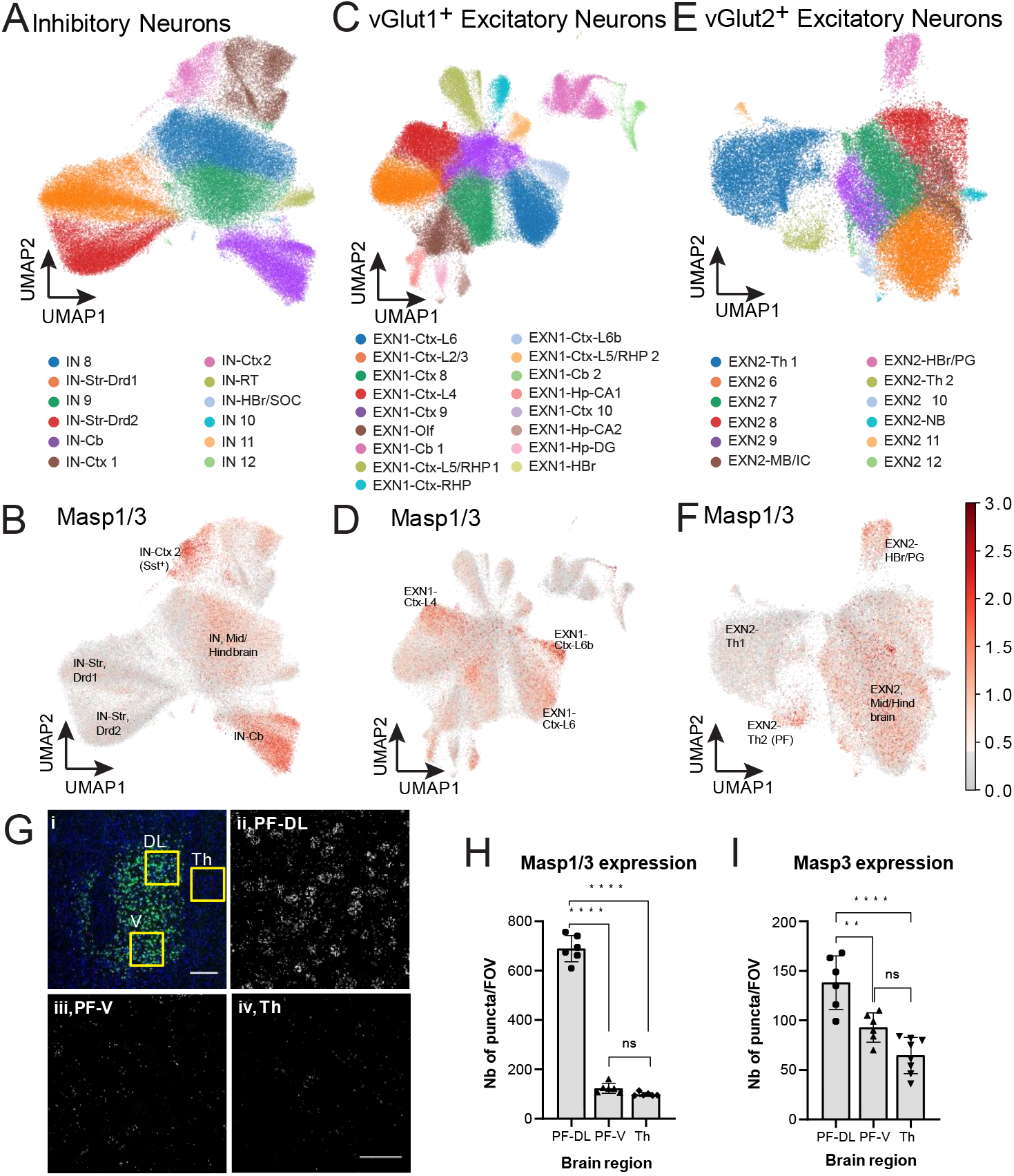
Spatial heterogeneity of Masp1/3 expression. **(A, B)** Masp1/3 expression by MERFISH projected onto a UMAP of secondary clustering of inhibitory neurons (A: UMAP of inhibitory neuron subclusters, reproduced from Figure S2E; B: projection of Masp1/3 expression on to inhibitory neuron subcluster UMAP). **(C, D)** Similar plots as in A and B for vGlut1^+^ excitatory neurons. C is reproduced from Figure S2A. **(E, F)** Similar plots as in A and B for vGlut2^+^ excitatory neurons. E is reproduced from Figure S2C. A Lists of cell type abbreviations is provided in Table S7. **(G)** Representative image of Masp1/3 (isoform non-specific) RNAscope in the parafascicular region and thalamus. i) Low resolution image of RNAscope of PF region marker Lypd6b. Highlighted regions indicate locations of images taken to evaluate Masp1/3 transcript abundance. Blue: DAPI nuclei; Green: Lypd6b. Scale bar: 200µm. ii-iv) representative images of Masp1/3 RNAscope puncta in dorsal lateral (PF-DL, ii), and ventral (PF-V, iii) side of the PF nucleus, and thalamus region adjacent to the PF nucleus (Th, iv). **(H)** Quantification of Masp1/3 puncta per FOV measured by RNAscope in respective sub-regions of the PF nucleus and thalamus. Scale bar: 50µm. n=6 mice, 1 FOV per mouse per region; One-way ANOVA followed by Tukey’s multiple comparison test; error bar shows standard deviation; ns: not significant; ****: p<0.0001. **(F)** Quantification of Masp3 puncta per FOV by isoform-specific RNAscope probe in sub-regions of the PF nucleus and adjacent thalamus region. n=3 mice, 2 FOV per mouse per region; One-way ANOVA followed by Tukey’s multiple comparison test; error bar shows standard deviation; ns: not significant; **: p<0.01; ****: p<0.0001.

### Masp3 has a profound effect on development and cognitive function

To study *in vivo* functions of Masp3, we turned to a Masp3 deficient mouse model (Gullipalli et al., 2023). In this model, exon 12 of the Masp1/3 gene was deleted via CRISPR/Cas9, which specifically inactivates Masp3, leaving Masp1 and MAp44 intact (Gullipalli et al., 2023). We confirmed the loss of AP activity in Masp3 homozygous knockout mice (Masp3 -/-) plasma by the rabbit red blood cell lysis assay (Gullipalli et al., 2023). The plasma AP activity in heterozygous Masp3 knockout mice (Masp3 +/-) was unaffected compared to wild type littermate controls (Masp3 +/+) (Figure S7A). We first observed that the birth rate of Masp3-/- was significantly lower than the expected Mendelian ratio: 20 out of 173 (11.6%, compared to expected 25% or 43 mice) female mice, and 23 out of 160 male mice (14.4%, compared to expected 25% or 40 mice) born from heterozygous breeding pairs were Masp3-/- (Table S5). Among those that survived, Masp3-/- mice had sustained lower body weight. On average, male Masp3-/- weighed 20% less than littermate control Masp3 +/+ males, and females weighed 14.5% less (Figure 4A). Together, these observations indicate a critical role of Masp3 in systemic development.

**Figure 4:**
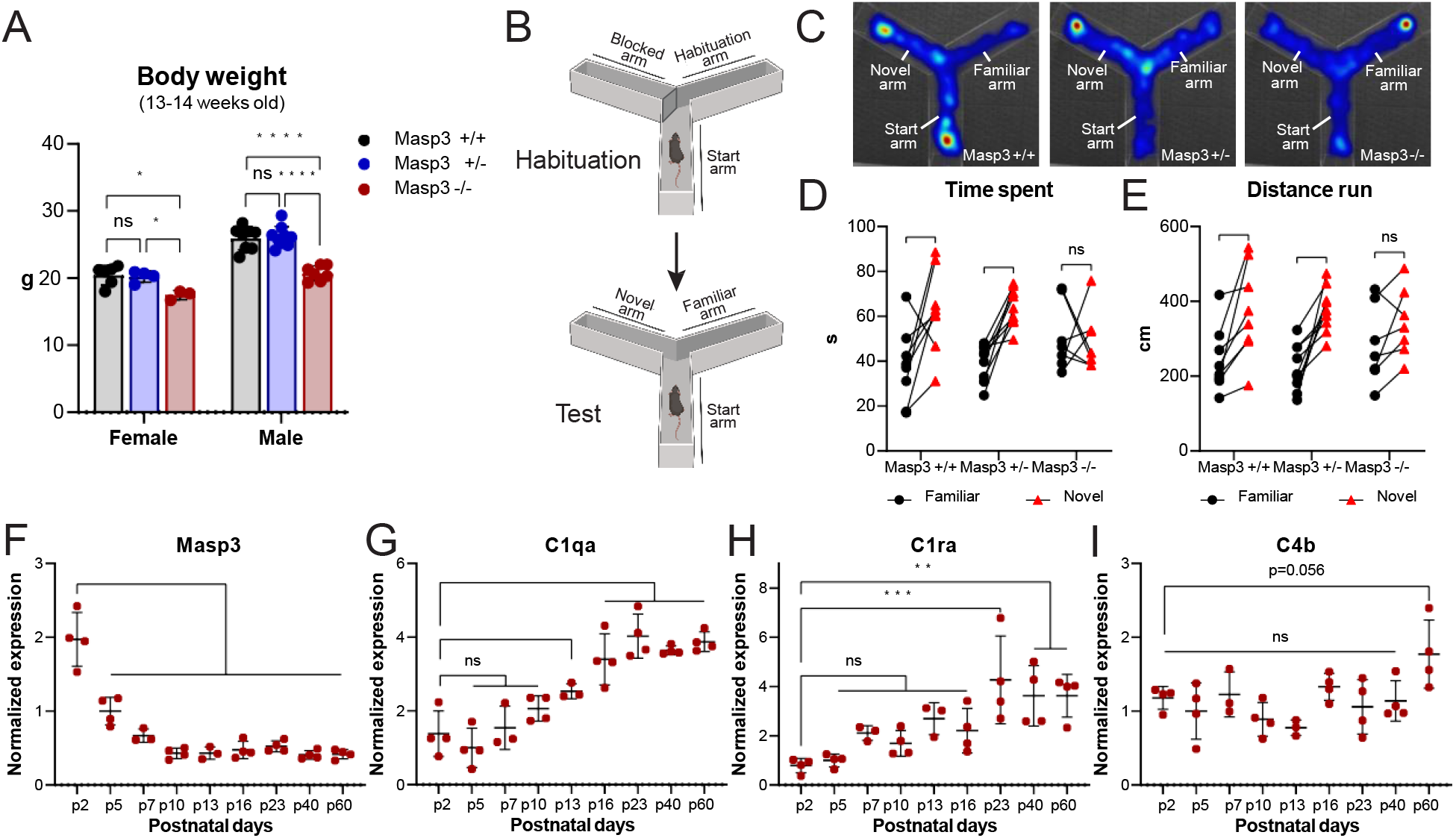
Masp3 deficiency resulted in cognitive deficits without affecting microglia dependent synapse pruning. **(A)** Body weight of 13- to 14-week-old adult Masp3+/+ (n=6 females, n=8 males), Masp3 +/- (n=4 females, n=9 males) and Masp3 -/- (n=3 females, n=7 males). Two-way ANOVA with Tukey’s multiple comparisons test; error bars represent standard deviation; ns: not significant; *: p<0.05; ****: p<0.0001. **(B)** Schematic representation of spatial novelty Y maze test for spatial working memory, created with BioRender.com. **(C)** Representative heatmap of spatial novelty Y-maze during test phase in which the left arm was novel, in Masp3 +/+ (left panel), Masp3 +/- (middle panel) and Masp3 -/- (right panel) mice. **(D-E)** Time spent (D) and distance run (E) in the familiar (black dots) vs. novel (red triangles) arm in spatial novelty Y-maze test for spatial working memory. Masp3 +/+: n=8; Masp3 +/-: n=9; Masp3 -/-: n=7; all male mice. Paired two-way ANOVA followed by Sidak’s test for multiple comparison; error bars represent standard deviation; ns: not significant; *: p<0.05; **: p<0.01; ***: p<0.001. **(F-I)** Time course qRT-PCR of Masp3 (F), C1qa (G), C1ra (H), and C4b (I) in mouse frontal cortex. n=4 mice per time point, except n=3 mice for p7 and p13. One-way ANOVA with Dunnett’s multiple comparison test; ns: not significant; *: p<0.05; **: p<0.01; ***: p<0.001; ****: p<0.0001.

Spatial and temporal expression patterns of Masp3 indicate it could be involved in multiple aspects of mouse behavior, assuming high expression implies functional importance. First, it is highly expressed in the entire developing cortex, rather than a particular region of the cortex. Therefore, it could regulate cortical functions such as cognition, anxiety, depression, or sensorimotor function. Second, it is highly expressed in the PG nuclei, an integral part of the cortico-ponto-cerebellar circuitry that is primarily known to regulate motor functions (Kratochwil et al., 2017). And third, it is highly expressed in the PF region, especially in the dorsal lateral side of the PF that projects to the caudate putamen. PF has been shown to regulate locomotion, motor learning, and depression depending on the projection (Zhang et al., 2022). Therefore, to test *in vivo* functions of Masp3, we performed a series of behavior tests focusing on cognition, anxiety, depression and motor function. A novelty Y maze test for spatial working memory was applied to assess cognitive function (illustrated in Figure 4B). Mice with intact cognitive function are expected to memorize the novel vs. familiar arm and are expected to exhibit a preference for the novel arm because of their exploratory nature. Both wild type control (Masp3 +/+, n=8, Figure 4C, left panel) and Masp3 +/- mice (n=9, Figure 4C, middle panel) showed such expected preference for previously unexplored novel arm, as measured by time spent (Figure 4D) and distance run in the novel vs. familiar arms (Figure 4E). However, the Masp3 -/- mice (n=7, Figure 4C, right panel) did not show such preference, indicating deficits in spatial working memory. Masp3 -/- mice are indistinguishable from WT and heterozygous mice for their level of anxiety (by light/dark box test, Figure S7B), and depression (by tail suspension (Figure S7D), and forced swim (Figure S7E) tests), and for their motor functions (by open field test for locomotion (Figure S7F), and rotarod test (Figure S7G)). These observations indicate that the predominant *in vivo* function of Masp3 resides in cortical development, especially in the establishment of neuronal circuitry related to cognition. On the contrary, its role in the PF and PG nuclei is likely non-essential.

To find the critical period of Masp3 expression during the course of postnatal development, we performed time-course quantitative RT-PCR on mouse brain frontal cortex samples (from postnatal day 2 (p2) to p60, n=3-4 per time point). The level of Masp3 kept declining after birth and reached its lowest expression at p10 (Figure 4F). On the contrary, CP components C1qa, C1ra both increased over postnatal development, and reached peak expression at p23 (Figures 4G-4H). Cortical C4b expression increased modestly over development as well, albeit statistically insignificant (Figure 4I).

Collectively, transcription data implied non redundant function of the AP and the CP during postnatal brain development. First, there were limited overlap of cell types that expressed components of the two pathways, with microglia (for C1q complex and C3), astrocytes (for C4b) and ependymal cells (for C3 and C4b) synthesize most of CP components, whereas neurons and neuronal progenitors are main producers of AP components Cfb, Cfd, Cfp and Masp1/3 (Figure 1M). Spatially, there was inverse correlation of spatial expression pattern of AP components with CP components (Figures 2H, S6A, S6C). And temporally, AP and CP are active during different brain development stages, with CP being more relevant in neuronal maturation much later in postnatal development, and Masp3 mediated AP being more important in early development.

## Discussion

Here we used MERFISH to produce a spatially resolved brain atlas of the complement system over postnatal development. With our approach, we found that the brain has a local supply of most complement components, whose expression exhibited remarkable temporal, cellular and spatial heterogeneity. Complement activation in the brain is strictly regulated. First, expression of serine proteases that are indispensable for complement cascade is maintained at very low level at steady state to keep minimal basal level complement activity. In addition, absence of C5 (or Hc) and C9 expression practically prohibited formation of MAC and subsequent lytic cell death in healthy brains, an aspect that is unique to the brain. Finally, complement regulatory proteins are abundantly synthesized both by neurons and non-neuronal cells in the brain to provide an additional level of complement regulation.

Although a role of the CP activation in normal brain development (Presumey et al., 2017; Schafer et al., 2012; Stephan et al., 2012; Stevens et al., 2007) and neurological diseases (Dalakas et al., 2020; Hong et al., 2016; Kjaeldgaard et al., 2018; Krance et al., 2021; Sekar et al., 2016; Shi et al., 2017; Watkins et al., 2016; Wilton et al., 2023; Yilmaz et al., 2021) have been widely reported, little is known about the LP and AP in the brain. Here, we provided transcriptional evidence arguing for an absence of the LP in the brain at steady state, another brain-specific aspect, based on the lack of expression LP serine proteases Masp1 and Masp2, and high-level expression of LP inhibitor, MAp44. However, it remains possible that the LP could be activated in aging and disease conditions, for example, in traumatic brain injury (De Blasio et al., 2017; Mercurio et al., 2020) and ischemic stroke in the brain (Fumagalli and De Simoni, 2016). On the other hand, our data revealed an unexpected presence of the AP in the brain during development, supported by the observation of abundant expression of Masp3 in early postnatal cortex that subsequently declined over development. Furthermore, the spatial distribution of Masp3 expression is highly correlated with that of Cfd, its direct downstream target that is essential to AP activation. Masp3 deficiency resulted in reduced birth rate, lower body weight and cognitive deficits, indicating a profound effect of Masp3 on systemic, as well as cognitive development. In humans, homozygous loss of function mutations in MASP3 causes 3MC syndrome, a rare autosomal recessive disorder characterized by delayed development, abnormal facial cranial structure, short stature, and intellectual disabilities (Atik et al., 2015; Pihl et al., 2017), further supporting a critical developmental role of MASP3. Although the most well-established molecular function of Masp3 is the proteolytic activation of Cfd, it may act on other substrates outside of the complement cascade (Cortesio and Jiang, 2006). To further confirm a role of the AP in brain development, Cfb and Cfd deficient mice should be examined.

Synapse pruning is the most well-characterized function of complement in the brain. During this process, activated complement component C3b deposits on synaptic terminals, which can then be recognized and engulfed by microglia through C3 receptor Itgam/Cd11b. Although this process has been mostly associated with the CP, the AP can also mediate proteolytic activation of C3 and therefore drive microglia-dependent synapse pruning. While this may be true, we observed inverse correlation of spatial and temporal expression patterns of genes involved in these two complement pathways, which indicates that the AP may have functions non redundant with the CP. Besides synapse pruning, less conventional roles of the complement pathway in neural development have been reported. Mice with C3 deficiency displayed reduced adult neurogenesis, both at basal level and in response to cerebral ischemia (Rahpeymai et al., 2006). Considering that Masp1/3 is highly expressed in neural progenitor cells, it is possible that AP could influence neurogenesis. Alternatively, complement could affect *in vitro* (Shinjyo et al., 2009) and *in vivo* (Gorelik et al., 2017b) neural progenitor migration. In fact, isoform non-specific ablation of the Masp1/3 gene by CRISPR resulted in aberrant embryonic neural migration (Gorelik et al., 2017b). Further study is needed to distinguish whether this phenotype is due to the loss of Masp1, Masp3 or MAp44.

The role of the AP activity in the brain may extend beyond development, and drive neurological disorders. AP activation has been observed in age-related macular degeneration (AMD), and AP components Cfb, Cfd and Cfh have been identified as risk factors for AMD (Edwards et al., 2005; Gold et al., 2006; Haines et al., 2005; Stanton et al., 2011). AP activation has also been shown in a mouse model for multiple sclerosis (MS), in which C3, but not C1q, localized to the site of neurodegeneration (Werneburg et al., 2020), indicating involvement of CP-independent complement pathway activation in MS pathogenesis. In addition, genetic ablation of complement factor B (Cfb), a key component downstream of Cfd that drives the AP activation, provided protection against demyelination, and these Cfb deficient mice had significantly reduced disease severity (Nataf et al., 2000). Hence, CP is likely not the sole player of the complement cascade in the central nervous system. We propose that deciphering differential involvement of each complement activation pathway in the pathogenesis of neurological diseases, although not commonly performed, is highly necessary for the development of effective complement therapies with improved safety profile.

## Methods

### Mice

All animal experiments were performed in compliance with the Institutional Animal Care and Use Committee at Harvard Medical School and Boston Children’s Hospital (protocol numbers IS00000748, IS00000111, IS00004762 and IS00002660), and with the guidelines of the Laboratory Animal Center at the NIH for the humane treatment of animals. C57Bl/6J mice (WT; The Jackson Laboratory, 000664) were bred and maintained in house. The Masp3 mutant mice were transferred from Song lab at University of Pennsylvania, School of Medicine, and expanded in house. Mice were maintained at the AAALAC-accredited animal facility at Harvard Medical School, with a 12-hour light-dark cycle and fed with standard chow diet and water ad libitum. 5-6 day old newborn pups and 8-14 week-old adult mice were used in for the study. 2-day old to 8-week old mice were used for the time-course gene expression study.

### MERFISH library target gene selection

We selected a panel of 249 genes composed of seven major categories: 1) complement related genes (a total of 52 genes, including 26 complement components, 13 known and putative complement receptors in the brain, and 13 complement regulators (Battin et al., 2019; Carroll, 2004; Hourcade, 2006; Kovacs et al., 2020; Krych et al., 1992; Ricklin et al., 2010; Sheehan et al., 1995; Tschopp et al., 1993; Zhou et al., 2023; Zipfel and Skerka, 2009); 2) brain cell type markers (2-3 well-established markers per brain cell type) (Codeluppi et al., 2018; Zeisel et al., 2018); 3) brain region markers selected based on empirical data and ABA reference ISH results; 4) genes related to neuronal function (including a comprehensive list of neurotransmitter synthesis, transports and receptors) and homeostasis selected based on empirical data; 5) astrocyte function related to neurodevelopment and disease conditions (Chung et al., 2013; Clarke and Barres, 2013; Zamanian et al., 2012); 6) microglia in neurodevelopment and disease conditions (Hammond et al., 2019; Keren-Shaul et al., 2017; Lehrman et al., 2018); 7) other genes implicated in neurodevelopmental disorders.

### MERFISH probe sequence design and probe library construction

MERFISH probe sequences were designed using a previously established protocol (Moffitt et al., 2016). Each target gene in our library was assigned a unique 24-bit binary barcode with hamming-distance 4, and hamming-weight 4. The Ensembl Genome Reference Consortium Mouse Build 38 (mm10) was used as reference transcript sequences. When a gene of interest has multiple isoforms, the predominantly expressed isoform was selected as the reference sequence. If multiple isoforms were expressed at a comparable level, probes were not penalized for targeting multiple isoforms. 80 distinct probes were designed for each target gene, though a few genes too short to support this number of probes had fewer. Each probe contains one 30-nt targeting sequence complementary to the target gene that has minimal homology to other genes. Overlap of the 30-nt complementary region between probes for the same target gene was allowed. To each targeting sequence, three 20-nt readout regions were concatenated, the sequence of which is complementary to the readout probe sequence whose associated bit in the barcode assigned to that gene contained a ‘1’. The probe sequences can be found in supplementary table 6. The template oligopool library was custom synthesized by Twist Biosciences, and amplified to generate the final probe library for hybridization following published protocol (Moffitt et al., 2016).

### MERFISH sample preparation, imaging and decoding of raw image data

Samples for MERFISH were prepared as previously described (Moffitt et al., 2016). Briefly, mice were euthanized with isoflurane overdose. Brains were quickly dissected and immediately frozen on dry ice and stored at -80°C until sectioning. Sagittal slices of 14-µm thickness were collected at 960±80µm, and 2040±80µm from the midline in p5 mice, and at 1000±100µm, and 2450±100µm from the midline in p60 mice. Brain slices were collected on poly-D-lysine coated silanized glass coverslips covered with orange fiducial beads that were prepared as previously described (Moffitt et al., 2016). Brain sections were allowed to air dry, and then fixed in 4% paraformaldehyde (PFA) for 10 minutes at room temperature, and washed three times with RNase-free PBS. Fixed brain slices were stored in 70% ethanol for at least 16 hours (up to 3 days) before hybridization. Encoding probe hybridization was performed at 37°C for 48 hours in a humidified chamber inside a covered petri dish as previously described (Moffitt et al., 2016). Samples were then washed, embedded in polyacrylamide gel, and treated with proteinase K as previously described (Moffitt et al., 2016). Fully prepared samples were stored in RNase-free 2X SSC in 4°C for less than 3 days before imaging. Images were collected on a custom microscope and fluidic system. Z stacks of 10-µm thickness at 1-µm step size were taken for each field of view. 12 cycles of two-color readout probe hybridization, imaging, fluorophore-cleavage, and wash were performed to measure signals for the 24-bit barcode scheme. The total imaging time for a full sagittal section was approximately 36-40 hours. To ensure optimal signal, buffers and readout probe solutions were replaced with fresh preparations after 6 cycles of imaging. Raw MERFISH imaging data was decoded using a previously established pipeline (Moffitt et al., 2016).

### MERFISH data analyses

In order to minimize sample-to-sample variation, only samples with comparable overall signal brightness and within 50% of variability in the number of total decoded RNA per cell were included in final analyses. Features with a volume above 100 µm^3^ were considered cells, those below the volume threshold were removed, as well as cells with zero total RNA counts. Data were normalized by cell volume and then scaled to bring the total number of RNA counts per cell to 250. RNA counts were then log transformed for cell type clustering with the Scanpy package (Wolf et al., 2018) following the general workflow. To correct for batch effect, the Harmony algorithm (Korsunsky et al., 2019) was applied after principal component analysis (PCA), and the subsequently generated corrected PCs were used for clustering. Primary clustering was performed using the Leiden algorithm (Traag et al., 2019) with kNN=30 and resolution=1. Neuronal cell types identified from primary clustering were regrouped into inhibitory neurons, vGlut1^+^ neurons and vGlut2^+^ excitatory neurons. Secondary clustering was then performed for each primary cell type with individually adjusted kNN and resolution to avoid over-clustering. Spatial analyses were performed with custom code in Python and is available upon request. Differentially expressed gene analysis was performed with a Benjamini-Hochberg correction for multiple comparisons.

### RNA expression analysis by ddPCR and qRT-PCR

Cortex and thalamus samples were dissected based on visible anatomical features after transcardiac perfusion with 20ml ice-cold RNase-free PBS. Brain tissue samples were lysed in the presence of TRIzol reagent (Ambion 15-596-018) with TissueLyser homogenizer (Qiagen 85600). RNA was isolated using phenol-chloroform extraction followed by purification with Zymo Direct-zol RNA Miniprep Kit (Zymo Research R2052). Genomic DNA removal and cDNA synthesis were performed with iScript gDNA clear cDNA synthesis kit (Bio-Rad 1725035) by following manufacturer’s instructions. ddPCR reactions were run with QX200™ ddPCR™ EvaGreen Supermix (Bio-Rad 186-4034). Each 20µl reaction mix contains 10µl EvaGreen Supermix, 0.5µl primer mix at 5µM each, and 9.5µl of cDNA. Droplets containing the reaction mixture was generated by the Bio-Rad microfluidic droplet generator, PCR amplified using the following cycle conditions: 95°C for 3min, 40 cycles of 95°C for 20 sec, 60°C for 20 sec and 72°C for 30sec, and then read with the Bio-Rad QX100 droplet reader. Data were analyzed using Bio-Rad QuantaSoft software v.1.7.4.0917. qRT-PCR was performed using iTaq^TM^ universal SYBR^®^ Green supermix system (Bio-Rad 1725124) and run on the Bio-Rad CFX96 real-time system with the same cycle conditions as in ddPCR. Signal was read at the end of each 60°C incubation period. Hprt1 was used as reference gene in both ddPCR and qRT-PCR. Primers for ddPCR and qRT-PCR were designed using the PrimerBank primer designing tool developed by Harvard Medical School/Massachusetts General Hospital. Hprt1 (fwd: GCGTCGTGATTAGCGATGATG; rev: CTCGAGCAAGTCTTTCAGTCC); Masp1 (fwd: AGTGCTCAAGAGAAGCCTGC; rev: AGCAGCTGTCAAAACCCAGT) (Hayashi et al., 2019); Masp3 (fwd: AGTGCTCAAGAGAAGCCTGC; rev: ACCCTCGATGTGTCTTCCAC) (Hayashi et al., 2019); MAp44 (fwd: CAAAGACCAAGTGCTCGTCA; rev: CTTCTCCAATTCGATCTCGC) (Hayashi et al., 2019); C1qa (fwd: AAAGGCAATCCAGGCAATATCA; rev: TGGTTCTGGTATGGACTCTCC); C1ra (fwd: GGCTCCATTTACCTCCCTCAG; rev: CACGTCAAACTGCCAGAAGAC); C4b (fwd: AGCCTGTTTCCAGCTCAAAG; rev: GTCCTAAGGCCTCACACCTG) .

### RNAscope^TM^ *in situ* hybridization

RNAscope^TM^ *in situ* hybridization was performed on fresh frozen brain sections of 14μm thickness. RNAscope^TM^ Fluorescent Multiplex Assay kit (ACDBio, 323130) was used. Samples were prepared by following manufacturer’s instructions. Probes: Mm-Lypd6b, channel 2 (ACD Bio 525851-C2); Mm-Masp1, channel 1 (isoform non-specific, ACDBio 525851-C1); Mm-Masp1-O1, channel 1 (Masp3-specific customized probe for NM_001359083.1, ACDBio 1176541-C1); Mm-Masp1-O2, channel 1 (Masp1-specific customized probe for NM_008555.3, ACDBio 1193891-C1); Mm-Masp1-O4, channel 1 (MAp44-specific customized probe for XM_006521829.5, ACDBio 201741-C1). Images were obtained using Olympus Fluoview FV1000 confocal microscope with 10X (for low resolution Lypd6b images) and 60X magnification objectives.

### Rabbit red blood cell hemolytic assay for alternative complement activation

Rabbit RBC hemolytic assay was performed as previously described (Gullipalli et al., 2023). Briefly, rabbit red blood cell (Complement technology, B300) was washed in GVBo (Ca^2+^ and Mg^2+^ free, Complement technology, B103) buffer and diluted to 10^9^/ml in the same GVBo buffer. Mouse EDTA plasma was diluted to 40% in GVB-Mg^2+^EGTA (5mM) buffer. Diluted plasma was incubated with 10µl of diluted rabbit RBC in a final volume of 50µl, and incubated at 37°C for 30 min in a water bath with mixing. For complete hemolysis control, 10µl diluted rabbit RBC was added to 40µl of H_2_O. 10mM EDTA was added at the end of incubation to stop the reaction. Cells were centrifuged at 3000rpm for 3min at 4°C. OD 405nm was read on the supernatant. Percent hemolysis was calculated as sample OD405nm divided by OD405 of RBC lysis in H_2_O.

### Behavioral tests

For all behavioral tests, mice were allowed to acclimate for at least 30 minutes.

#### Spatial novelty Y maze test

The experiment was run in a clear, Y-shaped 3-arm maze with a 120° angle between the arms. Each mouse was run twice in the Y-Maze, first for a 3 min habituation phase then for a 3 min test phase, with a delay of 2min from the end habituation phase and the beginning of test phase. The testing arena was cleaned after each trial. For each trial, the mouse was always introduced in the same arm, denoted as ‘starting arm’. For habituation phase, one arm was blocked off, which is randomized and balanced for genotypes. For test phase, the blockade was removed, the mouse was re-introduced and allowed to freely explore the entire testing arena. All trials were video recorded, and analyzed using EthoVision XT 17 software (Noldus). The time spent and distance run in the familiar and the novel arm was compared.

#### Light-dark box test

The experiment was run in ENV-520 open-field arena (Med Associates) that was equally divided into two chambers with a small opening in between. One chamber was under direct illumination (650 lux) and the other chamber was protected from light with a black opaque insertion. The test mouse was introduced to the center of the light chamber and was allowed to run freely in both chambers for 10 min. Its movement was tracked with infrared beams and all data were recorded and analyzed with the Activity Monitor software from Med Associates. The percentage of time spent and distance ran in the light chamber are reported.

#### Open field test for locomotion

The experiment was performed in a clear 27*27*20cm^3^ Plexiglas chamber from Med Associates. The test mouse was allowed to run freely in the test arena for 1 hour and tracked with infrared tracking system. Data were recorded and analyzed with the Activity Monitor software from Med associates. The total distance traveled (cm) was reported for general locomotor activity.

#### Rotarod test

The experiment was performed on a rotarod apparatus (Stoelting; Ugo Basile Apparatus) consists a texturized plastic roller flanked by large round dividers to isolate test mice. Mice are tested in groups of up to 5 mice. A habituation run with performed by letting the mice run on the rod rotating at 4rpm for 5min. Durin this step, mice were replaced back to run if they drop from the rotating rod. Following habituation, mice were reintroduced to the rotarod apparatus and tested with accelerating rod rotation speed (4-40rpm in 3min, maintained at 40rpm for a total of 5min). Mice were tested three times and the average latency to fall across three trials were reported. Mice were allowed to rest for at least 20min between trials.

#### Tail suspension test

In this experiment, a 4cm plastic thin tube was introduced to the test mouse’s tail as a climb stopper. A 5cm scotch tape was adhered and folded in half around the tip of the test mouse’s tail. The test mouse was hooked to the test arena and suspended upside down for 10min, without being able to touch the bottom of the test arena. Its movement was recorded, tracked and analyzed with the EthoVision XT 17 software (Noldus). The total immobile time was reported.

#### Forced swim test

This experiment was performed in 2000ml Pyrex beakers with a diameter of approximately 12cm, filled with 25±1°C water to the 1400ml mark. The test mouse was weighed before the start of the test and each trial duration was 6min. The test mouse’s movement was recorded, tracked and analyzed with the EthoVision XT 17 software (Noldus) and the total immobile time was reported.

### Statistics

Statistical analyses were performed in Python, GraphPad Prism 8.3.0. and Excel. All data in dot plots were presented as mean ± standard deviation. One-way or two-way ANOVA was applied for multiple group comparisons. Pair-wise comparisons were performed using appropriated multiple comparison tests. Detailed statistical methods used, p values, false discovery rate (FDR) were indicated in figure legends.

## Data availability

MERFISH data of this study will be deposited on datadryad.org upon publication. An interactive online platform: <u>Spatial and temporal atlas of the complement system in brain development (Zhang etal.) (harvard.edu)</u> will be made available to facilitate spatial data visualization, and to provide additional cell type, brain region and age dependent differential expression analyses.

## Acknowledgements

We would like to thank Dr. Aritra Bhattacherjee for discussion, advice and Dr. Chao Zhang for support in data analyses. We would like to thank Elisabeth Elicot for administrative support and Diana Pascual for mouse colony management.

## Author contributions

M.C.C. conceived the project; M.C.C., J.R.M. supervised the project; Y.Z., B.C., J.R.M., M.C.C. designed the experiments; Y.Z. designed and performed MERFISH experiments and analyzed data; B.W. constructed the MERFISH instrument, performed user training, and provided essential reagent preparation; T.L., S.C., L.G. constructed the MERFISH data visualization website; Y.Z. performed qRT-PCR, ddPCR, RNAscope; A.R. performed rabbit red blood cell hemolytic assay; Y.Z., Y.G. performed mouse behavior tests; M.M. performed mouse genotyping; W.S. generated the Masp3-/- mouse model; Y.Z., J.R.M., M.C.C. wrote the manuscript.

## Declaration of interests

J.R.M is a co-founder of, stake-holder in, and advisor for Vizgen, Inc. J.R.M. is an inventor on patents associated with MERFISH applied for on his behalf by Harvard University and Boston Children’s Hospital. J.R.M.’s interests were reviewed and are managed by Boston Children’s Hospital in accordance with their conflict-of-interest policies.

## Supplementary figure legends

**Figure S1:**
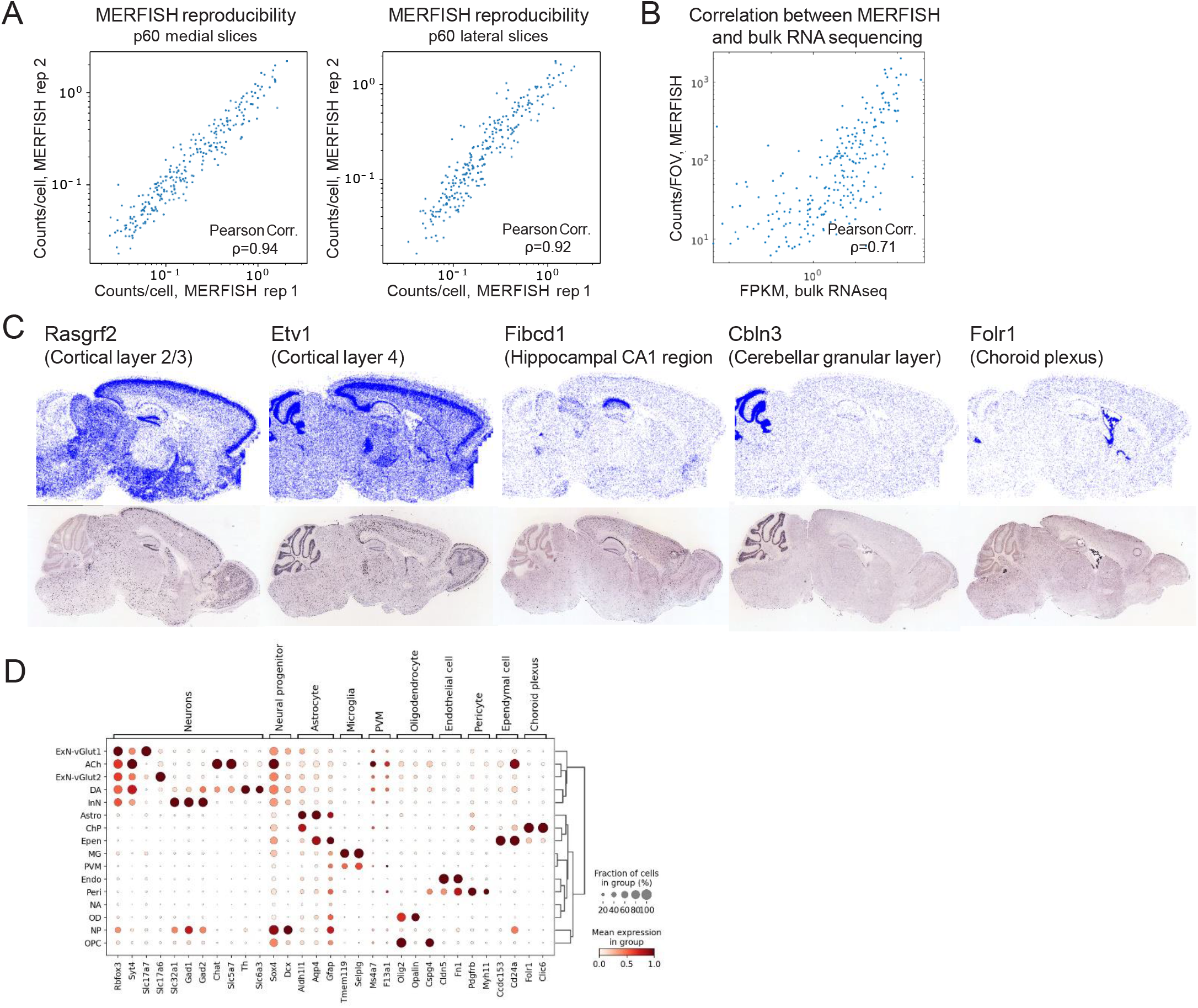
MERFISH reproducibility and consistency with brain bulk RNA sequencing and with Allen Brain Atlas in situ hybridization database. (A) Scatterplot of RNA counts per cell of individual genes measured by MERFISH in two independent adult mouse brain medial (left panel) and lateral (right panel) sagittal slices. **(B)** Scatterplot of RNA counts per field of view (FOV) of individual genes measured by MERFISH (p60, medial slice) and FPKM measured by bulk RNA sequencing of brain cortex. **(C)** Representative plots of spatial distribution of a subset of brain region markers included in MERFISH target gene library (top panels) and reference in-situ hybridization (ISH) images from Allen Brain Atlas (ABA, bottom panels). Left to right: Rasgrf2: cortical layer 2-3 marker; Etv1: cortical layer 4 marker; Fibcd1: Hippocampal CA1 region marker; Cbln3: cerebellar granular layer marker; Folr1: choroid plexus marker. **(D)** Dot plot of major brain cell type marker expression in primary cell clusters identified in p60 mouse brains by MERFISH. Color intensity represents average expression, and size of the dot represents percentage of cells expressing each gene.

**Figure S2:**
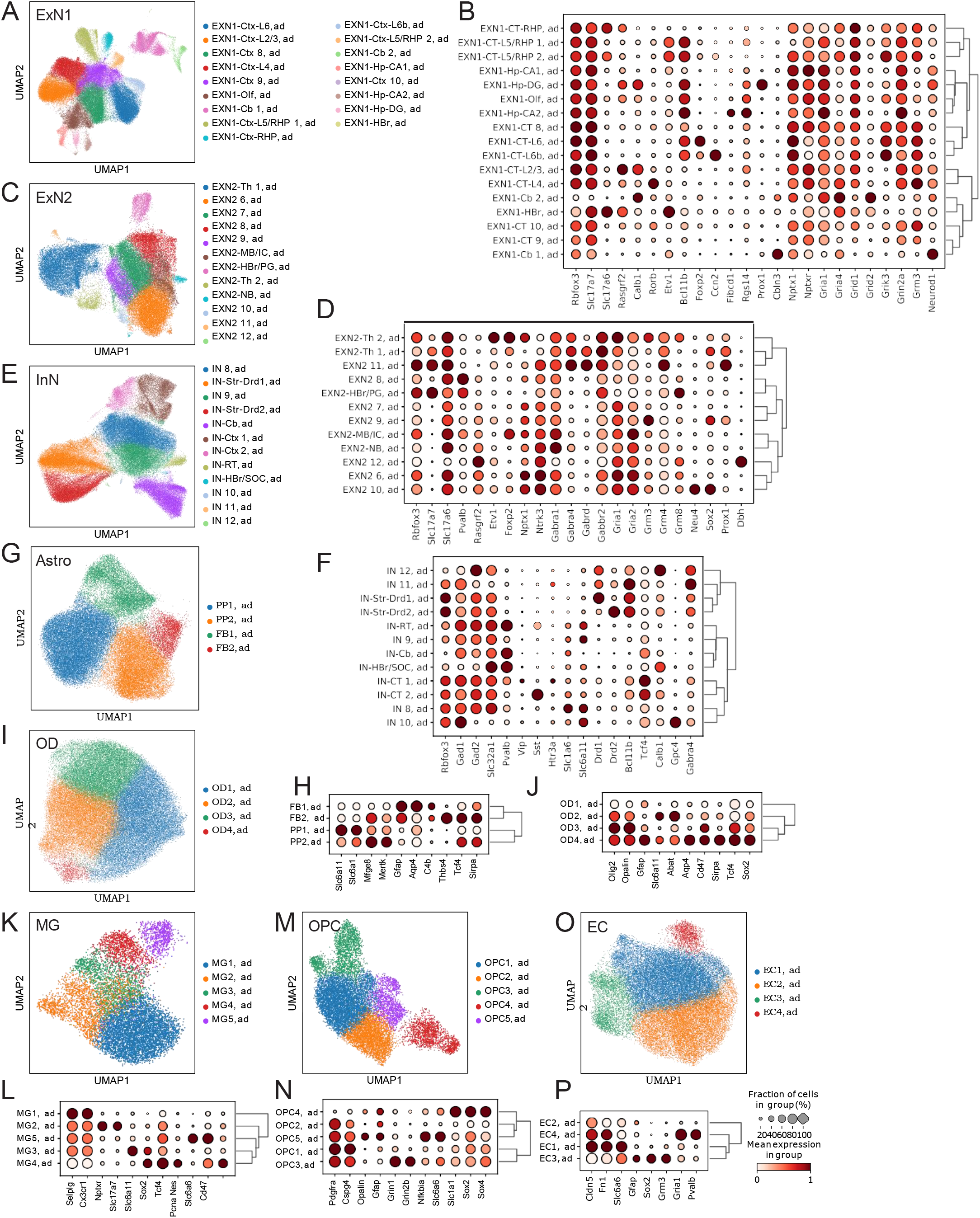
Secondary clustering of p60 samples. UMAP (A, C, E, G, I, K, M, O) and dot plot of top differentially expressed genes (B, D, F, H, J, L, N, P) of vGlut1^+^ (ExN1, A,B), and vGlut2^+^ (ExN2, C, D) excitatory neurons, inhibitory neurons (InN, E, F), astrocytes (Astro, G, H), oligodendrocytes (OD, I, J), microglia (MG, K, L), oligodendrocyte progenitor cells (OPC, M, N) and endothelial cells (EC, O, P). For dot plots, color intensity represents average expression, and size of the dot represents percentage of cells expressing each gene. A, C and E are reused in figure 3C, 3E and 3A respectively for clarity of the main figure.

**Figure S3:**
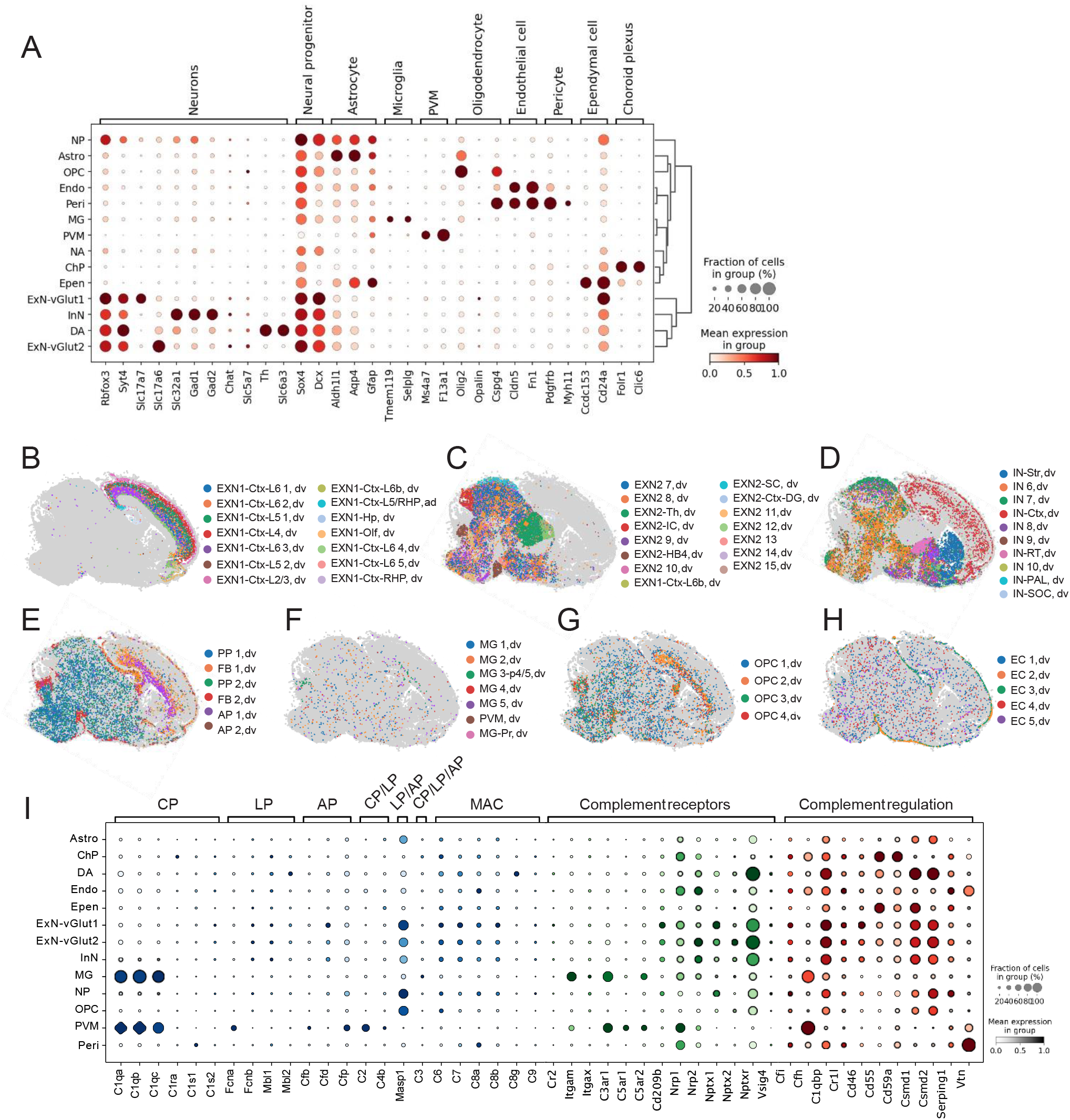
Complement expression in p5 juvenile mouse brain. **(A)** Dot plot of major brain cell type marker expression in primary cell clusters identified in p5 mouse brains by MERFISH. **(B-H)** Spatial localization of secondary cell clusters of vGlut1^+^ (B) and vGlut2^+^ (C) excitatory neurons, inhibitory neurons (D), astrocyte (E), microglia (F), oligodendrocyte progenitor cells (G) and endothelial cells (H). **(I)** Dot plot of complement expression by cell type. In dot plots (C and M), color intensity corresponds to average gene expression, dot size corresponds to the percentage of cells expressing each gene.

**Figure S4:**
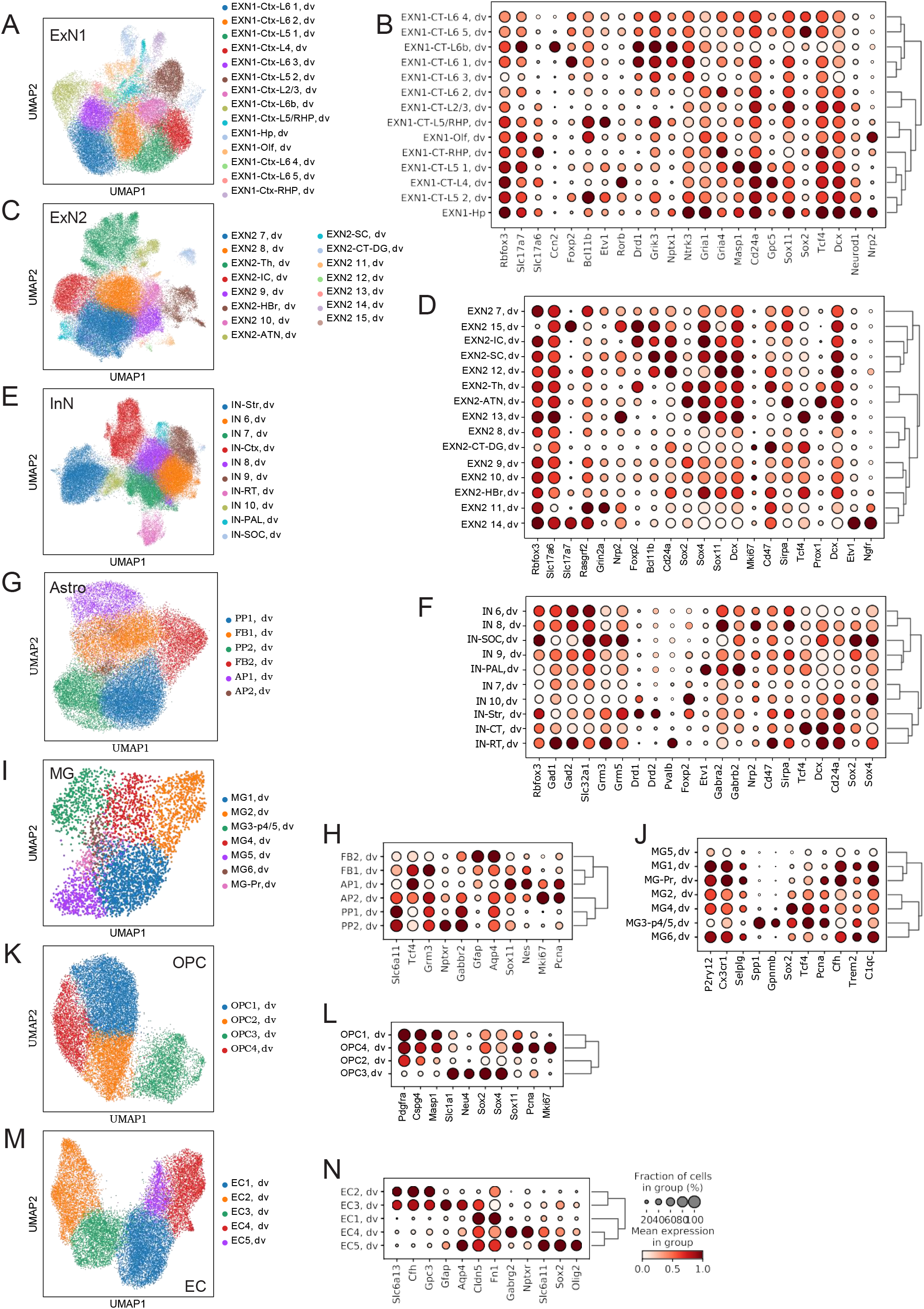
Secondary clustering of p5 samples. UMAP (A, C, E, G, I, K, M) and dot plot of top differentially expressed genes (B, D, F, H, J, L, N) of vGlut1^+^ (ExN1, A,B), and vGlut2^+^ (ExN2, C, D) excitatory neurons, inhibitory neurons (InN, E, F), astrocytes (Astro, G, H), microglia (MG, I, J), oligodendrocyte progenitor cells (OPC, K, L) and endothelial cells (EC, M, N). For dot plots, color intensity represents average expression, and size of the dot represents percentage of cells expressing each gene.

**Figure S5:**
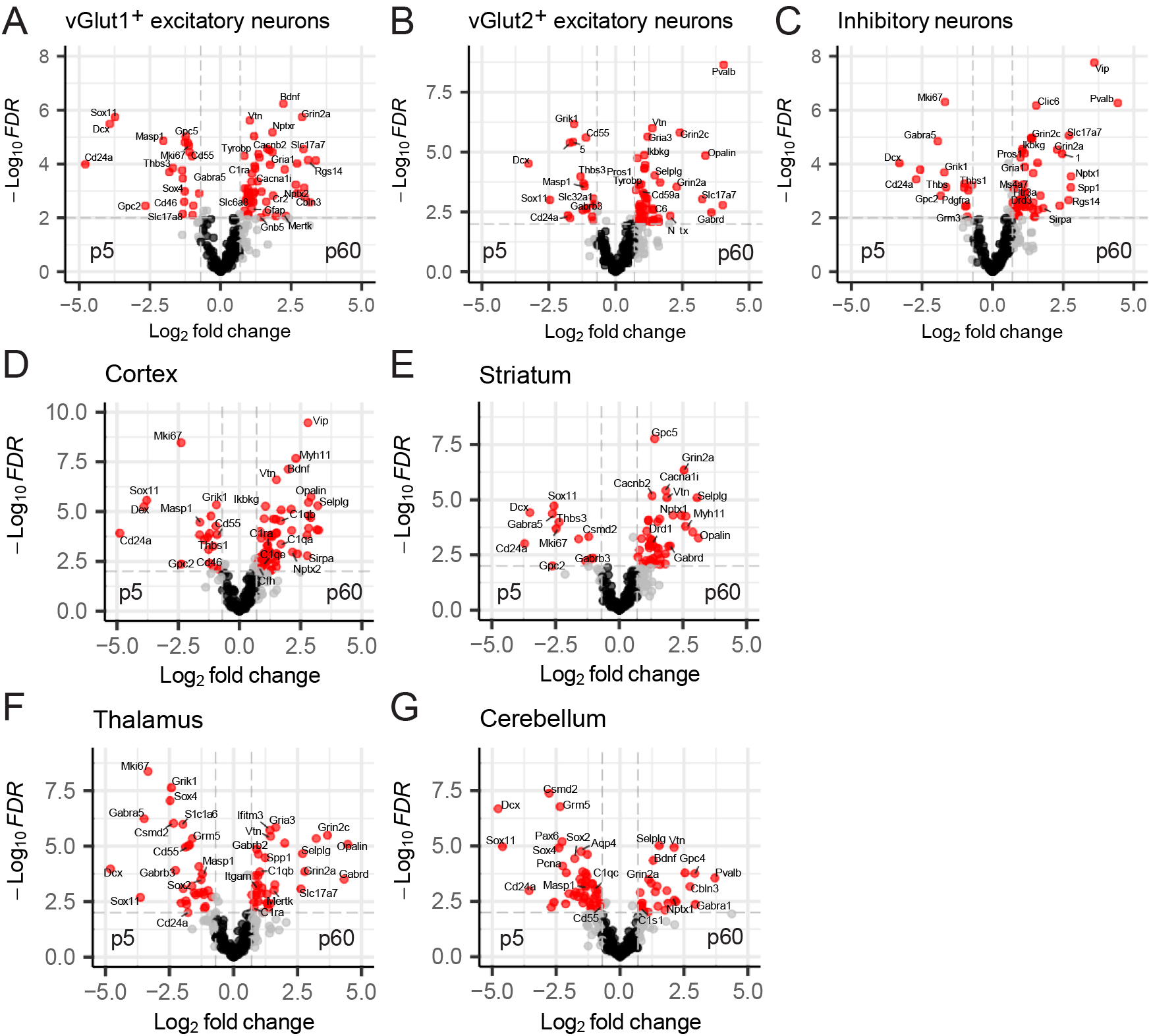
Developmentally regulated differential gene expression by cell type and brain region. Volcano plot showing cell-type specific **(A-C)** and brain-region specific **(D-G)** differentially expressed genes in p5 vs. p60 mouse brains detected by MERFISH. (A) vGlut1^+^ excitatory neurons; (B) vGlut2^+^ excitatory neurons; (C) inhibitory neurons; (D) all cell types in the cortex; (E) all cell types in the striatum; (F) all cell types in the thalamus and (G) all cell types in the cerebellum. Genes with over 1.6-fold difference between p5 and p60, and with false discovery rate (FDR, calculated by Benjamini-Hochberg test for multiple comparisons) less than 0.01 are highlighted in red.

**Figure S6:**
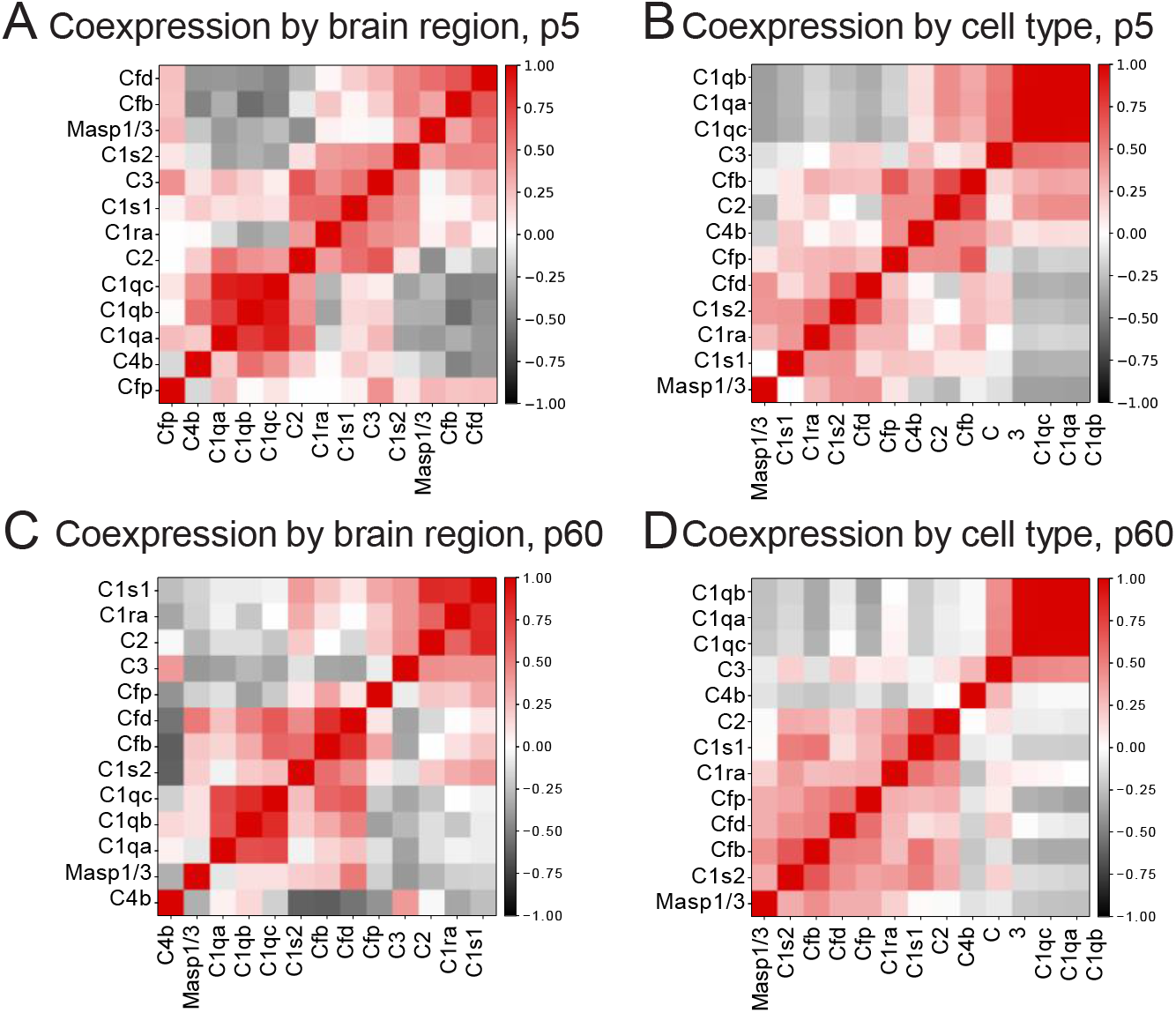
Co-expression pattern among components of all three complement activation pathways. Heatmap of pairwise Pearson correlation coefficient of complement component expression by brain region (A, C), and by cell type (B, D). A, B: from p5 juvenile mice; C, D: from p60 adult mice. Complement genes detected below MERFISH detection threshold in both age groups were not taken into account in the analysis. For co-expression pattern by brain region analyses, average gene expression was calculated for each common brain region shared between the two age groups were used; for co-expression pattern by cell type, average gene expression was calculated for each refined cell cluster.

**Figure S7:**
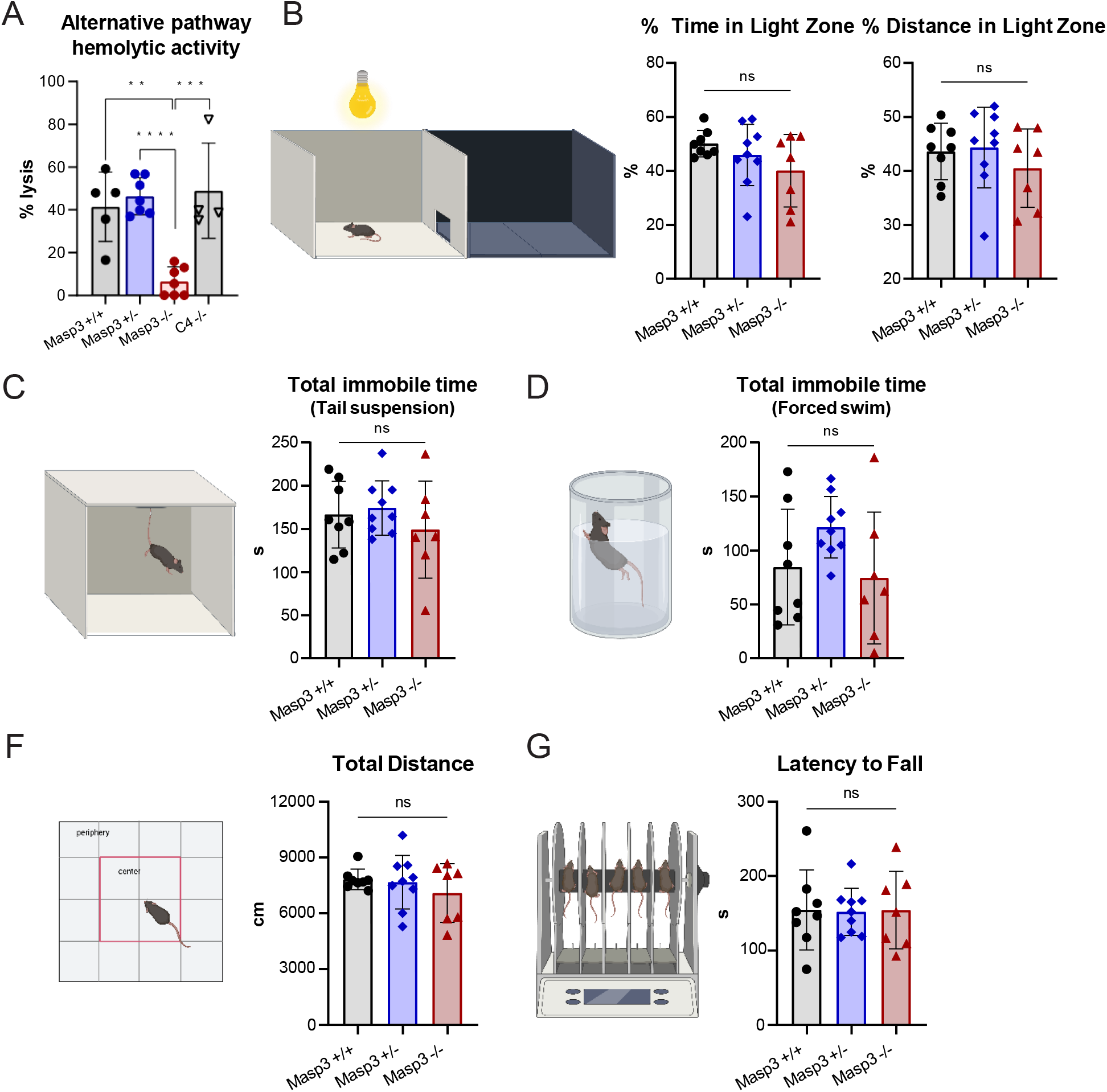
Masp3 deficiency doesn’t affect anxiety, depression and locomotion. (A) Rabbit red blood cell hemolytic assay with Masp3 +/+ (n=5), Masp3 +/- (n=7), Masp3 -/- (n=7) and C4b-/- (n=4) EDTA plasma. Percent hemolysis was reported by considering hemolysis in water as 100%. Error bars represent standard deviation. One-way ANOVA followed by Tukey’s test for multiple comparison. **: p<0.01; ***: p<0.001; ****: p<0.0001. **(B)** Light/dark box text for the level of anxiety. Percent time spent, and percent distance run in the light zone was reported. **(C, D)** Tail suspension test (C) and forced swim test (D) for depression-like behavior. Total immobile time was reported. **(E)** Total ambulatory distance run in open field test for locomotion. **(F)** Latency to fall in rotarod test for motor coordination. Average of three tests are plotted for each mouse. All behavior tests were run on male animals with Masp3 +/+: n=8; Masp3 +/-: n=9; Masp3 -/-: n=7. Error bars represent standard deviation. One-way ANOVA followed by Tukey’s test for multiple comparison was applied. ns: not significant. No significant abnormality was detected in Masp3-/- mice in the above behavioral tests. Illustrations of behavioral test setup were created with BioRender.com.

## Notes

### Competing Interest Statement

Jeffrey R. Moffitt is a cofounder of, stake-holder in, and advisor for Vizgen, Inc. J.R.M is an inventor on patents associated with MERFISH applied for on his behalf by Harvard University and Boston Children's Hospital. J.R.M's interests were reviewed and are managed by Boston Children's Hospital in accordance with their conflict-of-interest policies.

